# The hippocampus as a small-world cognitive map

**DOI:** 10.64898/2026.02.07.704615

**Authors:** Jason Z. Kim, James P. Sethna, Itai Cohen, Weinan Sun

## Abstract

When a mouse perceives a hawk’s shadow, it may have only seconds to decide where to run, yet the safest refuge is often neither visible nor nearby. To survive, it must search its cognitive map quickly enough to choose among many possibilities, and accurately enough to avoid dead ends and hazards along the way, all while what counts as “safe” changes as paths, refuges, and threats shift with time. This scenario highlights a core design problem: cognitive maps must preserve fine local structure for reliable action, yet remain globally searchable so that distant, useful solutions can be found efficiently in both space and time. The hippocampus is thought to support such maps, with population activity representing world states and their transitions—yet how these maps are structured to solve this design problem is not well understood. Here, we used a novel geometry-aware autoencoder to model the structure of the cognitive map from longitudinal calcium imaging from thousands of hippocampal CA1 neurons in mice learning a memory-guided navigation task. We discovered that the hippocampal population code achieves both local fidelity and global searchability through **small-world network structure** in the space of neural representations that leverages two complementary mechanisms. At the population level, helical (rotation-plus-drift) dynamics of neural representations relative to past experience build new maps that preserve local information about nearby positions in space and time while remaining distinguishable from earlier representations. At the cellular level, neurons with coordinated multi-field activity create sparse, long-range shortcuts between distant representations. During synchronous population events in immobility, decoded activity often jumps to distant states in space, time, and task conditions, suggesting these shortcuts are engaged during offline processing. This functional organization, with implications for both neuroscience and artificial intelligence, sheds light on how hippocampal representations may be optimized for a fundamental challenge faced by intelligent systems: efficiently searching through accurate internal models of the world.

## 1 Introduction

A fundamental challenge faced by intelligent agents, from foraging rodents to chess grandmasters, is how to search an internal model of the world to guide behavior. Consider a rat in a maze-like burrow network. To recover food cached weeks earlier, it must use memory to navigate among tunnels, exits, and chambers without physically checking each branch. Or consider a person planning a route through an unfamiliar city, mentally simulating turns and landmarks before taking a single step. This capacity for *mental navigation*, the ability to efficiently search within an internal world model rather than through exhaustive physical exploration [1–3], is arguably what separates sophisticated cognition from simple reactive behavior.

The notion of a *cognitive map*, an internal representation that captures the relational structure of the environment and the organism’s state within it, provides a conceptual framework for understanding how such mental navigation might be achieved [1]. An effective cognitive map must satisfy two fundamental requirements. First, it must accurately represent the structure of the environment, preserving the relationships between states so that mental traversal reflects real-world transitions. Second, for mental navigation to be useful, the map must support efficient search that scales gracefully as environments grow in size and complexity. Without such efficiency, mental simulation would not be fundamentally faster than physical exploration.

The hippocampus offers a natural place to search for such organizational principles. It has long been implicated as the neural substrate supporting cognitive maps [4]. The landmark discovery of place cells, neurons that selectively fire at specific spatial locations, provided the first concrete evidence for neural encoding of an internal map [5]. Subsequent work has revealed that the hippocampus codes not only for spatial location but for a wide variety of experiences: temporal intervals [6], social identity [7, 8], conceptual categories [9, 10], and task-relevant abstract variables [11–13]. These findings support the broader conception of the hippocampus as encoding a general-purpose cognitive map [14].

Yet a critical question remains: **how is this map organized to support both accurate and efficient mental navigation?** Foundational work mapping single-neuron activity onto pre-selected variables such as position [5, 15] established what the hippocampus encodes, but leaves open how the population code is organized to support computation. Focusing on individual neurons provides only a partial view of the collective code [16]. To model the collective code researchers have turned to nonlinear dimensionality reduction techniques such as UMAP [17] and isomap [18] that produce low-dimensional embeddings of neural data, where the embedding coordinates are meant to reflect systematic changes in population activity associated with encoded variables. These embeddings, however, often lack the geometric fidelity needed to characterize representational structure: small displacements in the embedding can correspond to large, irregular changes in the predicted neural activity [19, 20], making it difficult to quantitatively study the organization of the map and how it supports navigation.

Here, we address this issue by using Γ-autoencoder (Γ-AE), a novel technique that uses differential geometry to build manifold models that more faithfully preserve distances, angles, and directions when mapping between the latent embedding coordinates and neural population activity [20]. We apply this method to longitudinal calcium imaging from thousands of hippocampal CA1 neurons tracked across weeks in mice learning a memory-guided navigation task [13].

We discover that the hippocampal cognitive map has **small-world network structure** [21] in representational space, combining dense local neighborhoods with a limited set of long-range “worm-hole” links between distant states. We trace this organization to two complementary features of the population code. First, helical representational dynamics, in which population activity associated with external spatial coordinates systematically rotates through state space while drifting across experience, maintain local continuity while keeping representations from different days distinguishable. Second, generalized state fields, in which coordinated multi-field activity recurs across otherwise distant neighborhoods, create sparse long-range shortcuts. Together, these mechanisms suggest an organizational principle by which hippocampal representations can support efficient search through accurate internal world models.

## 2 Quantitative model manifold of hippocampal cognitive maps

To study the collective representation of the mouse hippocampus, we use calcium imaging data of hippocampal CA1 neurons from a recently published study [13]. Briefly, the activity of thousands of neurons expressing the calcium indicator GCaMP6f was tracked across multiple days for 11 mice as they learned to navigate a virtual reality maze (Fig. 1a). The maze consisted of looped linear tracks with three zones: indicator, near reward 1 (R1), and far reward 2 (R2). Between trials a 2 s dark teleportation period was used to reset the track. For each trial, one of two indicator markers were randomly selected, which determined the location where the mouse needed to lick in order to receive a water reward (two-alternative cue-delay-choice (2ACDC) task) (Fig. 1b). Cranial windows were implanted to image the neural activity of CA1 neurons in the dorsal hippocampus (Fig. 1c).

**Figure 1:**
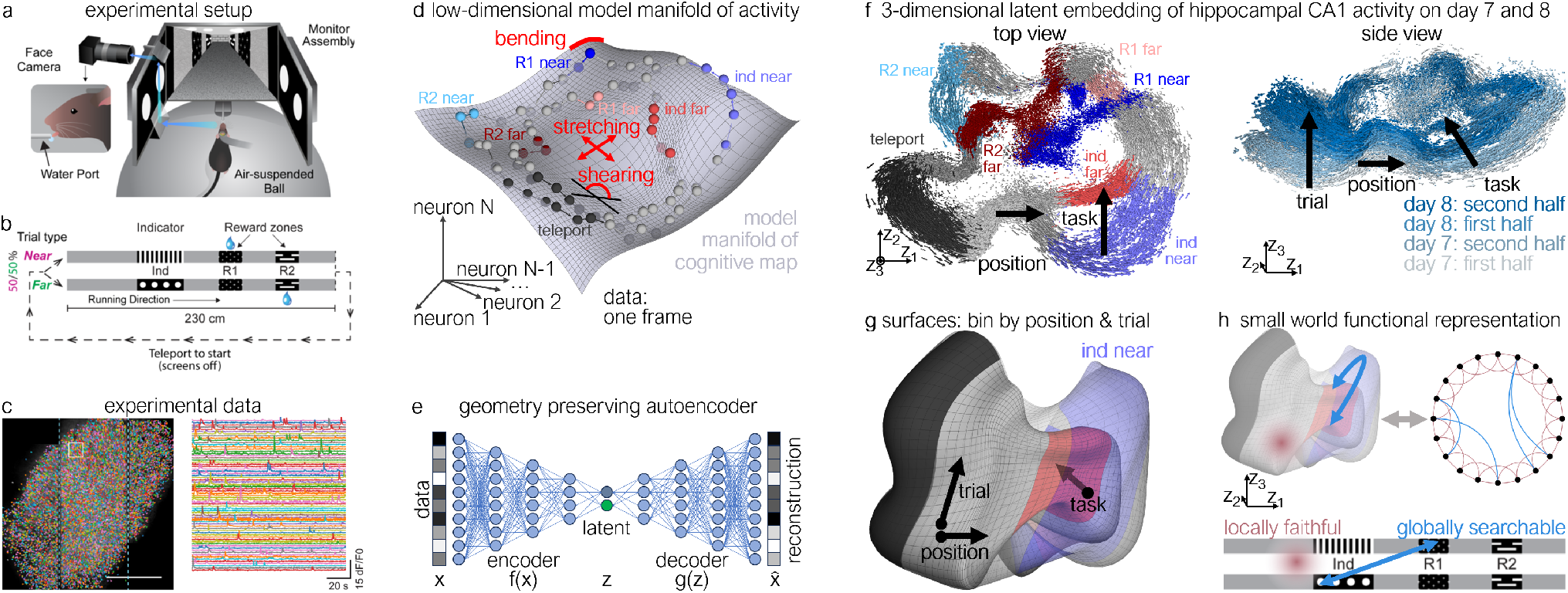
Constructing a quantitative model manifold of hippocampal cognitive maps. (**a**-**c**) Experimental setup (panels adapted from Sun et al. [13]). (**a**) Schematic of the experimental setup comprising a mouse navigating a virtual reality maze atop an air-suspended ball with simultaneous calcium imaging of hippocampal CA1 neurons. (**b**) Diagram of the linear maze with different indicator markers and corresponding reward locations. (**c**) Image ROI (left) and time traces of activity for a subset of neurons (right). (**d**) Schematic of constructing a low-dimensional geometric manifold of high-dimensional neuron activity, with sources of geometric distortion of bending, stretching, and shearing shown in red. (**e**) Architecture of the geometry preserving autoencoder that regularizes these distortions. (**f**) Embedding of neural activity on days 7 and 8 in a 3-dimensional latent space colored by track marker (left), and trial number (right), showing the three latent space dimensions encode for position, task condition, and trial. (**g**) Surfaces of the embedded data binned by position and trial for each task condition. (**h**) An effective cognitive map is locally faithful and globally searchable.

We use these longitudinally-tracked data across all days for each animal to model the cognitive map as a low-dimensional geometric manifold. Each imaging frame of *N* neuron activations defines a point in an *N* -dimensional space of population activity (Fig. 1d). We model these data using a novel geometry-regularized autoencoder (Γ-AE) [20] that constructs a low-dimensional geometric manifold directly in the high-dimensional space of neuron activity, mathematically suppressing geometric stretch, shear, and bend distortions of the manifold (Fig. 1d,e, see Methods). The latent variables define a coordinate system for the manifold where changes along the latent space faithfully correspond to movement along the manifold in data space. Thus, by explicitly preserving local distances, angles, and directions, Γ-AE yields a quantitative and interpretable model of population activity, where each dimension captures a different activity pattern.

We discover that the cognitive map codes for the position along the track, the task condition, and the trial number. Specifically, we plot the projection of neural activity onto a 3-dimensional latent space learned by Γ-AE for a representative mouse (Fig. 1f, see Methods) and color the points by track marker and trial number. As the mouse moves along the track, its neural activity locally evolves along one direction in the latent space, producing a loop for each task condition (Fig. 1f left). The two loops remain separate along a second local dimension from the indicator all the way to the end of R2, even when only the indicator is visually distinct. We further observe trials where the cognitive map model switches loops partway through a trial, often during error trials (licking at the wrong reward zone) or when the mouse is being inattentive (running without licking, see Supplemental Fig. 5), indicating that the second local dimension corresponds to the task condition in which the mouse believes itself to be. Crucially, a third prominent local dimension emerges across trials, along which neural activity systematically drifts in correspondence with accumulated experience and the passage of time (Fig. 1f right). This drift is a population-level manifestation of representational drift, the gradual reorganization of neural activity patterns encoding the same environment over time, and a well-known hallmark of hippocampal coding [22–25]. Because the majority of the mouse’s experienced mental state exists on two distinct task conditions, we define the model of the mouse’s *experience* on the cognitive map as the two surfaces formed by averaging the coordinates of the latent space data according to discrete bins of position (3cm) and time (20 trials, Fig. 1g, see Methods). In the following sections, we use our model to discover that the cognitive map has a small world functional representation that is both locally faithful and globally searchable (Fig. 1h).

## 3 Manifold rotation & drift contextualize local representation on unique past experience

These surfaces of experience provide a quantitative lens into how the mouse’s local representation of the maze evolves across trials (Fig. 2a). At each point on the surface, we compare two local directions: the position direction, defined by the change in population activity as the animal moves downstream along the track within a trial, and the trial direction, defined by the change in population activity when the animal returns to the same track location on later trials. A natural null expectation is that these directions should be orthogonal, since position and accumulated experience are distinct variables. In that case, trial number could still separate successive position rings in population space, but it would not rotate them relative to one another: corresponding track locations would remain phase-aligned across trials. Instead, we find almost everywhere on the surface that the trial direction forms a strikingly acute angle, *θ*, with the position direction (Fig. 2a). Geometrically, the grid line obtained by holding the track position fixed and moving to later trial bins does not rise straight away from the position ring; it leans toward the downstream side of the ring. In other words, when the animal returns to position *x* on a later trial, the population activity is shifted slightly toward the pattern that used to represent the downstream position *x*+Δ. The same relative shift can be viewed in the opposite reference frame: if we fix a neural activity pattern and ask where along the track it is expressed, that pattern is now elicited earlier along the track, matching classical place-field backshift phenomenon [26, 27]. Thus, viewed by following the same physical location across trials, the representation advances in phase around the neural activity loop. In the data, this phase advance is accompanied by drift into new activity patterns, so the representation both rotates relative to position and remains distinguishable across trials. We refer to this coupled phase advance (rotation) and drift as *helical dynamics*. Repeating this analysis across all animals (see Methods), we find that the distribution of *θ* (*n* = 11 mice; Fig. 8) is significantly acute, with a mean of *θ* ≈ 77.2° ± 2.7° (*p <* 0.001). Scaling this angle by the lengths of the position and trial vectors yields the phase advance of the representation along the track coordinate from one trial to the next, which is significantly positive across animals, averaging ∼0.43 mm/trial, or roughly 5 cm per session (Fig. 9, *p <* 0.001).

**Figure 2:**
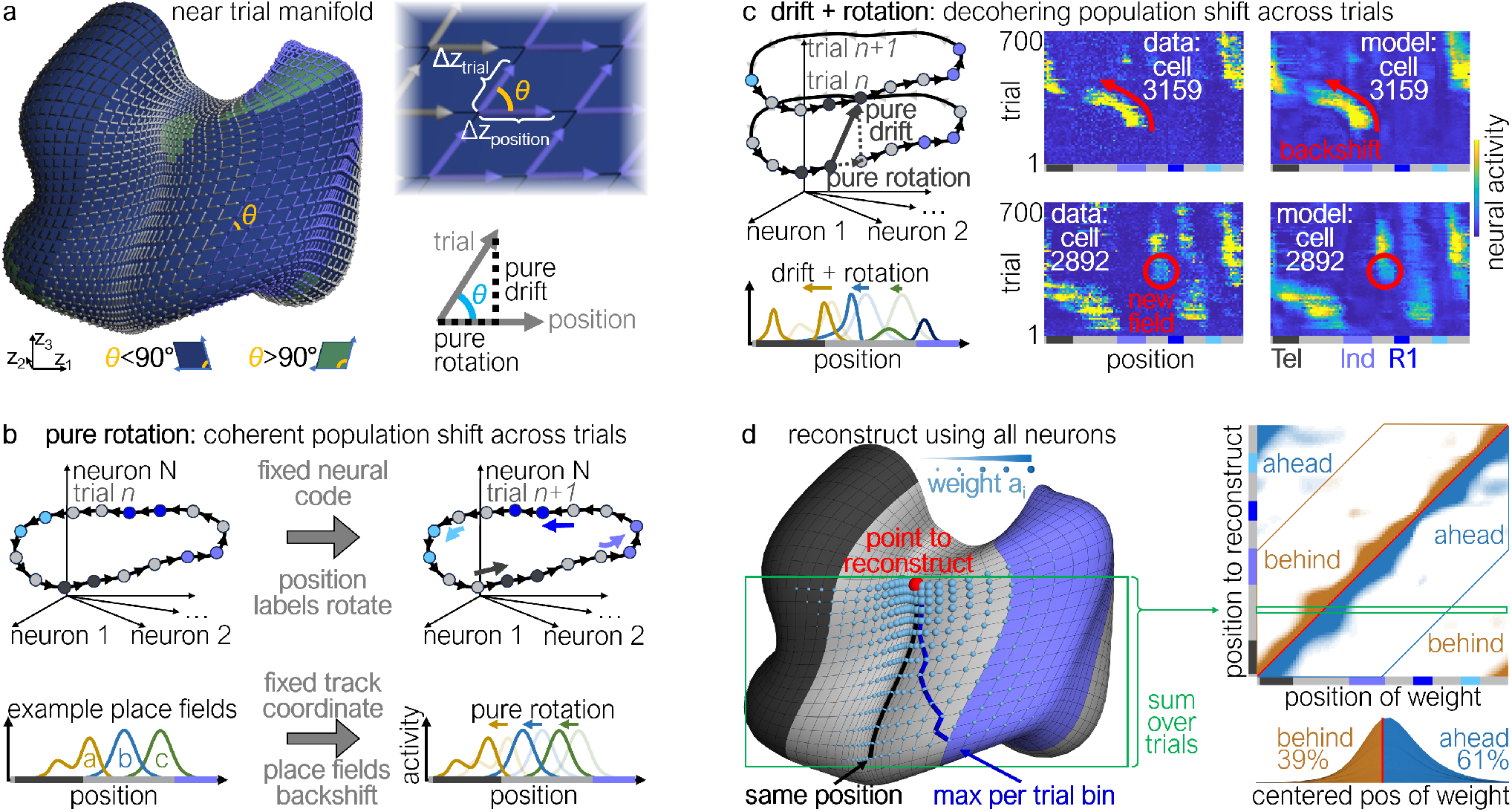
Manifold rotation and drift contextualize current state relative to unique past experiences. (**a**) Position- and trial-binned manifolds for the near task condition (left), with each bin colored by whether the angle between the vectors along position and trial is acute (blue) or obtuse (green), and zoomed-in view of one manifold section and schematic showing the angle, and the decomposition of the evolution across trials into a pure rotation and pure drift (right). (**b**) Schematic of the trajectory in the space of neuron activity for two consecutive trials undergoing pure rotation (top), associated with a consistent backshift of place fields (bottom). (**c**) Same schematic for two consecutive trials undergoing rotation and drift (top left), associated with a common component of backshift and a differential change in the place fields, respectively (bottom left), and example activity across position and trials for two neurons from the data and decoded from the 3D latent space model (right). (**d**) Compositionally reconstructing a decoded point on the surface using decoded bins from past trials and other positions results in weights that are skewed towards further positions on the track on earlier trials (left). Summing the weights over trials, and repeating this process for all positions on the same trial bin, we find that the weights are consistently skewed towards future positions on the track (right).

To build intuition for these helical dynamics, consider the activity patterns associated with one task condition during one trial as a ring in population-activity space (Fig. 2b). If we read out fictive neurons *a, b*, and *c* along this ring as functions of external position, a pure relative rotation would translate their activity profiles toward earlier positions on the track, matching classical place-field backshift [26, 27]. Equivalently, from the fixed-position view used above, the same operation means that each location recruits a population pattern that previously belonged to a slightly downstream location. However, pure rotation only reuses the same set of population activity patterns, and therefore cannot by itself explain why successive trial bins separate from one another on the surface. The observed helical progression must also include a component that carries the representation into new population-activity patterns. We therefore decompose the across-trial change into a rotational component, given by its projection along the position direction, and an orthogonal component that we call pure drift (Fig. 2c, top left). We associate both components with representational drift [22–25]: rotation preserves and rephases structure from prior experience, whereas drift keeps later experiences distinguishable from earlier ones. At the single-neuron level, pure drift can arise from changes in place-field shape, the formation or disappearance of fields, or even different rates of field backshift across neurons (Fig. 2c, bottom left). We find that all of these sources are observed in individual neuron activity and captured by the model (Fig. 2c, right, see Fig. 6,10 for more examples).

The fact that both rotation and drift exist implies that there is information from adjacent positions in previous trials that is preserved in (rotation), and also distinct from (drift) the current representation, thereby allowing past experiences to provide an explicit context for current representations. In this sense, helical dynamics bring the experienced future into the present: the representation of a current state is pulled toward activity patterns that previously occurred at upcoming states, while drift prevents it from collapsing onto those past patterns completely. As this process accumulates across trials, the current representation can carry a progressively broader trace of states that tend to follow it. To quantify this context, we use a compositional approach by reconstructing the activity pattern at a specific state on our surface using a non-negative weighted sum of the patterns at prior states, and use the weights as measures of the prior context. We find that the strongest weights begin centered around the point to reconstruct for the same trial bin, and shift towards further positions along the track for earlier trials (Fig. 2d, left). Summing the weights across trial bins and repeating this procedure by reconstructing each position, we find that the vast majority of the weights lie immediately ahead and behind that position (Fig. 2d, top right), with a significant skew ahead (Fig. 2d, bottom right). This skew towards upcoming states is reminiscent of successor representations previously observed in hippocampal CA1 neurons [28, 29], but generalizes to the population code which we thus term *generalized condensed representation* (GCR) (see Fig. 13 for a minimal geometric model of how successor-representation–like single-neuron tuning produces this rotation-plus-drift structure). Intuitively, GCR means that each state is represented together with a “compressed” imprint of its local neighborhood, forming dense local transition structure in the cognitive map that can preserve accurate relationships among nearby states through time.

## 4 Small-World Representational Architecture of the Hippocampal Cognitive Map

How, then, does the cognitive map connect distant representations? The manifold model gives us a way to ask this question at the level of single neurons. Rather than measuring a neuron’s tuning only as a function of a pre-selected variable such as position on one day, we can now use the decoder to view its activity as a field over the learned representational manifold, spanning position, task condition, and accumulated experience. When viewed over the full manifold, individual neurons form localized islands of activity that occupy restricted neighborhoods of position, task condition, and experience time, extending the notion of place fields beyond physical position alone (e.g. cell 2034, Fig. 3a). We call these localized regions generalized state fields (GSFs, see Methods), and define statistically significant local maxima within them as GSF peaks. Many neurons contain more than one such localized region (Fig. 3b), with different neurons occupying different fractions of the latent-space volume (Fig. 3b, right). Thus, multi-region GSFs generalize multi-field place cells to the full modeled cognitive map: a single neuron can participate in multiple, separated neighborhoods of representational space. These repeated localized fields provide a candidate mechanism for linking distant regions of the cognitive map.

**Figure 3:**
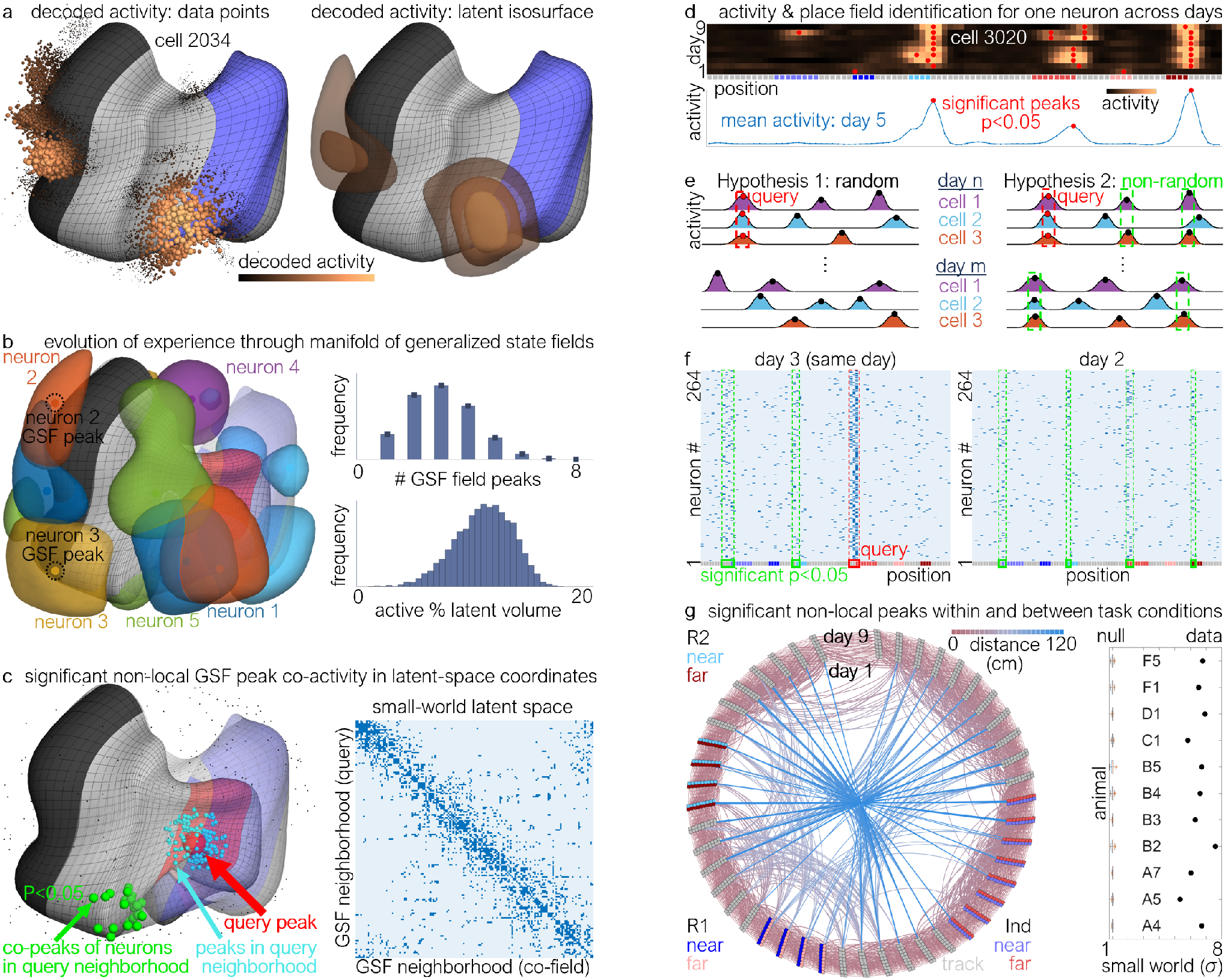
Small-world architecture of the cognitive map from the model and data. (**a**) Model activity of a single cell (2034) decoded from the embedded data points in the latent space (left), and isosurfaces of that same cell’s activity in the full latent space volume (right). (**b**) Isosurfaces of volumes with high single-cell activity for multiple neurons that tile the latent space we term Generalized State Fields (GSFs, left), and the distribution of GSF peak number (top right) and fraction of the latent space occupied by GSFs (bottom right). (**c**) Example of one query GSF peak (left, red), its nearest *k* = 200 neighbors (teal), the other GSF peaks of the query and neighbor neurons (co-peaks, black and green dots), and the co-peaks that are significantly clustered (green). Adjacency matrix of the small-world graph with connections between GSF neighborhoods when the significant co-fields of one neighborhood overlap with another neighborhood (right). (**d**) Rate map across positions and days for one neuron with marked peaks (red, top), and example trace from one day (bottom). (**e**) Schematic of one hypothesis of peak organization where co-peaks of neurons that have peaks at the query are randomly dispersed within and between days (left), *versus* another hypothesis where co-peaks are systematically organized (right). (**f**) Raster plot of neuron peaks for neurons that fire at the query, and with significant peak counts at different positions on the same day (left) and earlier day (right). (**g**) Connectogram where the nodes are position and day bins, and edges exist between the query bin and the statistically significant co-peak (left), and observed *versus* degree-sequence preserving null distribution of small-world coefficient for all animals (right).

We find that a significant number of neurons are simultaneously active at multiple latent space coordinates (Fig. 3c, left), suggesting that the neural code may have additional structure that results from the co-firing of neurons. Specifically, we consider the peaks of all GSFs in the latent space, query them one by one (Fig. 3c, left red), identify the neurons associated with the *K* = 200 nearest neighboring GSF peaks (Fig. 3c, left teal), and see whether the remaining GSF peaks of those neurons are (Fig. 3c, left green) or are not (Fig. 3c, left black) significantly clustered (see Methods). We observe that many neurons with peaks in the neighborhood of the query also have significantly clustered co-active peaks at *different* and *distant* latent space coordinates (Fig. 3c, right), which sometimes correspond to different positions, task conditions, and days. Viewed as a graph over latent-space neighborhoods, this combination of strong local clustering and a sparse set of long-range links yields a **small-world** organization [21] of the learned representational space (Fig. 3c, right; see Methods).

To test whether long-range links exist not just in the model of the cognitive map, but also between distant regions of physical space and time, we return to the experimental data to determine if there are a statistically significant number of neurons that co-fire at multiple positions on the track. It is well known that a place cell can fire at multiple positions over an environment [30](Fig. 3d, see Methods). Based on our model predictions (Fig. 3c), we hypothesize that neurons that co-fire at a specific query position, rather than firing randomly across positions and days (Fig. 3e, Hypothesis 1), will systematically co-fire at specific track positions and perhaps even other days (Fig. 3e, Hypothesis 2). Indeed, for neurons firing at one specific query position, we observe that a statistically significant subset of them also fire at different regions and on different days (Fig. 3f, see Methods, and Fig. 11 for raw unfiltered activity across single trials for these neurons). We repeat this analysis using each position and day bin as the query and compute the bins that contain a statistically significant number of neurons that co-fire. We plot a graph of bins across all positions, task conditions, and days, where edges exist from each query bin to other bins that have a significant number of co-firing neurons, and find that approximately 1% of the links in the graph are associated with long-range connections (Fig. 3g, left). We identify such sparse links in the graphs of all animals, and further find that the vast majority of these long-range connections are not captured using the same procedure on a smaller subsample of neurons (Fig. 12), consistent with the idea that detecting this sparse structure benefits from large-scale longitudinal recordings that reliably track thousands of neurons over days. Thus, in accordance with the model prediction, we find significant nonlocal co-activity of neurons at multiple and distant locations. Quantifying this graph against degree-preserving rewired null networks confirmed significant small-world organization in all animals (Fig. 3g, right; *p <* 0.001 for all 11 animals; see Methods).

## 5 Small-World Cognitive Map is Leveraged During Mental Navigation

To test whether this small-world architecture in the functional representation is engaged during behavior, we analyze neural activity when the animal stops and pauses in the task, a behavioral state often associated with hippocampal offline processing and memory-related reactivation rather than active task engagement [31]. Specifically, we examine activity patterns called *synchronized calcium events (SCEs)* that occur when the mouse is stationary (speed = 0) and is not licking. SCEs recorded with calcium imaging have been shown to co-occur with electrophysiologically measured sharp-wave ripples (SWRs) [32, 33], fast population events implicated in memory consolidation, reactivation, and memory-guided computation [31]. To decode whether these SCEs contain some component of a previously experienced position and task condition, we again compositionally reconstruct each SCE as a non-negative weighted sum of the neuron activity patterns associated with each position and day bin (Fig. 4a), and only keep weights that are statistically significant under a shuffle test (see Methods). We find that some SCEs decode to nearby states (Fig. 4b, red boxes) close to the animal’s actual position and day (black box). These SCEs that have locally decoded states resemble sequential re-activation patterns often observed in SWRs [31]. We also observe SCEs that decode remotely and non-contiguously from the animal’s position on the same day (Fig. 4c), resembling recent findings of super-diffusive SWR [34, 35]. Finally, we also observe SCEs that decode completely remotely in position, task condition, and across several days (Fig. 4d,e), consistent with recent findings that offline ensemble reactivation can link memories formed days apart [36].

**Figure 4:**
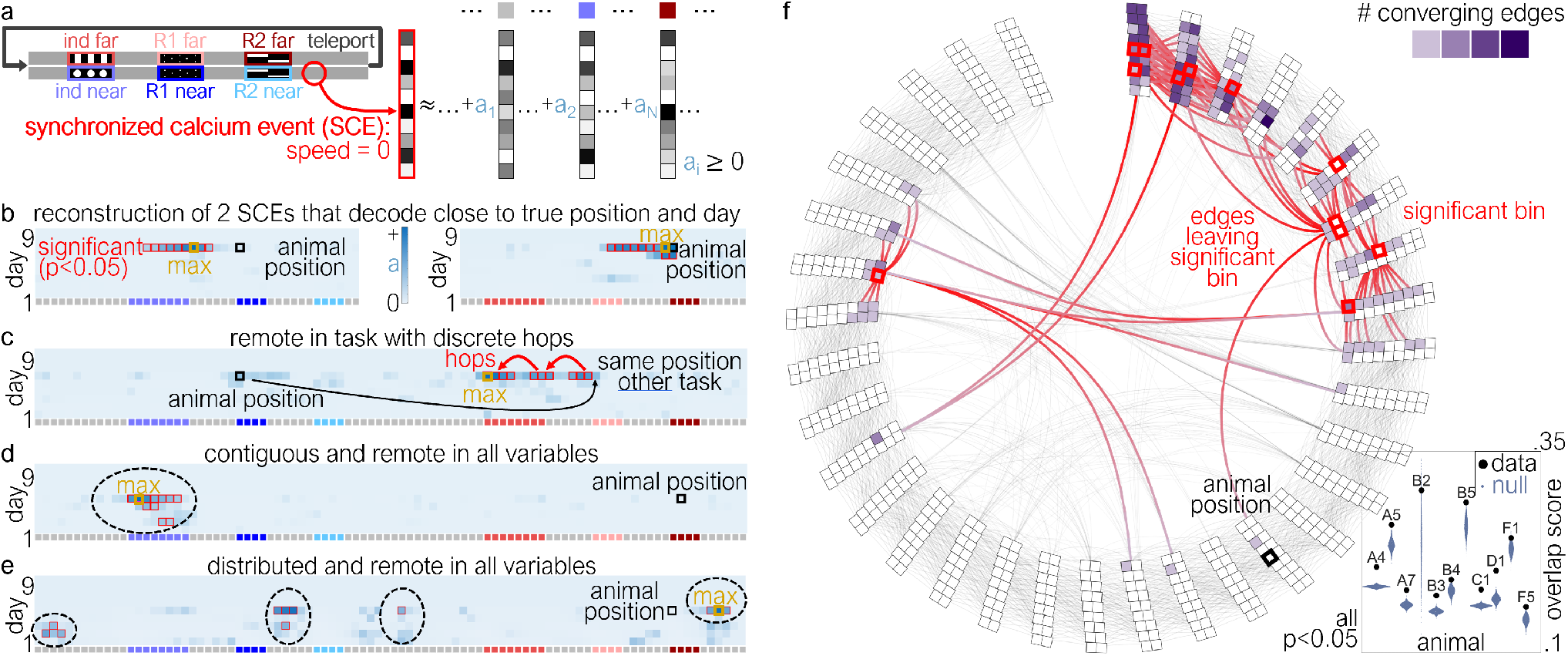
Accurate and efficient maps are leveraged during navigation. (**a**) Schematic of compositionally decoding synchronized calcium events (SCEs) which are activity patterns when the mouse speed was zero and was not licking. (**b**) Examples of significant reconstruction coefficients (red boxes) from two SCEs that decode close to the animal’s true position (black box) and day. Other examples of significant SCE coefficients that decode (**c**) remotely to the other task but contiguously in position and day, (**d**) contiguously and remotely in all variables, (**e**) and distributed and remote in all variables. (**f**) Statistically significant overlap between significant SCE reconstruction coefficients and one step down the small-world graph (left), and observed *versus* null distribution of the overlap score for all animals (right).

We next asked whether the decoded SCEs are organized by the small-world connectivity identified above (Fig. 3g), rather than simply activating arbitrary sets of states. If SCEs engage this graph, then the states decoded within a single event should be connected to one another by edges in the small-world map more often than expected by chance. To test this, for each SCE we took the bins with significant decoding weights (Fig. 4f, left, red boxes), followed the small-world graph one step from those bins, and measured how strongly this one-step graph neighborhood overlapped with the SCE’s decoded bins (purple squares; see Methods). For SCEs containing four or more significant decoding weights, this overlap score was significantly higher than a shuffle null in every animal (Fig. 4f, right). Thus, the same small-world organization inferred from representational geometry and coordinated multi-field activity is also reflected in pause-associated population events: SCEs can remain anchored to local map structure while also accessing remote states through sparse long-range links. Together, these results suggest that the hippocampal cognitive map is not only organized as a small-world network in representational space, but that this organization is expressed during pause-associated population events, providing a mechanism by which locally accurate maps can support nonlocal access to distant states.

## 6 Discussion

In this study, we use Γ-AE [20] to build a low-dimensional and geometrically faithful model of the cognitive map in hippocampal CA1 population activity across weeks of learning [13], thereby capturing how task states are quantitatively related in population activity space. We find that this map has small-world organization [21, 37]: strong local clustering preserves fine structure, and a few long-range shortcuts connect otherwise distant states. Crucially, this small-world structure is not in the anatomical wiring, but in the functional space of representations: how population activity states relate to each other. We trace the local component to helical (rotation-plus-drift) representational dynamics across experience, and the long-range component to coordinated multi-field structure in generalized state fields that tile the cognitive map. Immobility-associated synchronous population events decode to remote states that align with these bridges, linking representational topology to offline population dynamics.

### 6.1 Local structure from helical population dynamics

The dense local neighborhoods of the cognitive map arise from helical dynamics in which the mapping between external track position and population activity progressively shifts in phase relative to past experience, while the activity patterns themselves drift into new, distinguishable regions of population space in a coherent low-dimensional way. Individual neurons exhibit a wide range of experience-dependent changes, including place-field backshift, skew, drift, and shifts in reliability [22, 23, 26, 27, 38–47]. Our framework provides a population-level lens: heterogeneous single-neuron changes collectively reveal coherent low-dimensional helical structure.

This structure connects naturally to predictive-map ideas such as the successor representation (SR), in which each state is represented as a vector encoding expected future state occupancies relative to the external environment [28, 29]. The SR provides a powerful framework for understanding how hippocampal representations support flexible planning, and recent work has shown that SR-like structure can emerge from biologically plausible learning rules [48–51]. In our data, helical dynamics produce a related but more general organization, which we call the generalized condensed representation (GCR). Through rotation, the representation of a current state is pulled toward activity patterns that previously represented nearby upcoming states; through drift, this predictive overlap is embedded in new activity patterns rather than simply reusing the past. Thus, each point on the manifold carries a compressed trace of its local representational neighborhood across experience, including position, task condition, time, and any other variable present in the population code. In this sense, GCR is a population-level, geometry-based generalization of successor-like coding: rotation provides predictive local overlap, while drift expands capacity and separates experiences across days and task states. Helical dynamics therefore provides a locally navigable scaffold for the cognitive map, but efficient search across distant states requires shortcuts.

### 6.2 Long-range bridges from generalized state fields

The sparse long-range bridges that complete the small-world architecture arise from generalized state fields. Localized “fields” have long been used to describe hippocampal coding of physical and task-relevant abstract variables [5, 6, 12]. Our GSFs generalize this idea by defining fields directly in the geometry of the representational space: localized regions on the manifold of population activity spanning any variable represented in the population code. GSFs describe localized structure without specifying relevant variables *a priori*. Thus, multi-field firing commonly observed in hippocampal neurons [30, 52] becomes a computational feature by linking states that are distant in physical or representation space *via* coordination across neurons: cells that co-fire in one neighborhood co-fire at specific distant neighborhoods.

At first glance, these coordinated non-local secondary fields differ from prior work arguing that multi-field structure is largely independent across cells. In particular, Rich *et al*. reported that in a very large, primarily homogeneous linear environment, the number and locations of CA1 place fields were well explained by a stochastic model with minimal evidence for structured co-activation across distinct fields [30]. Subsequent studies have nevertheless highlighted conditions under which hippocampal coding exhibits additional structure, including experience-dependent coordination among neurons with *path-equivalent* (repeated) fields across related locations in environments with repeating elements [53], non-random *anatomical* clustering of CA1 neurons with similar field locations (often near salient task features and goals) [54], context-dependent structure in CA1 coactivity dynamics that carries information beyond single-cell tuning [55], and stronger correlated “cell-assembly” organization in CA3 consistent with recurrent-network mechanisms that can stabilize and bind distributed representations [56]. Thus, multi-field structure may become more salient with state-/task-dependence. Importantly, our chance-level controls explicitly account for the empirical inhomogeneity of peak density by estimating, and subsequently standardizing by, multinomial null probabilities (Eq. 10; Methods §7.11).

We clarify that these long-range bridges do not appear as topological shortcuts in the smooth, continuous Γ-AE manifold itself. The manifold provides a locally faithful geometry of population activity, whereas the shortcut structure emerges as a sparse functional overlay from coordinated firing: only certain neurons that participate in one representational neighborhood also participate together in distant neighborhoods of the latent space. This sparsity is likely a feature rather than a limitation. During local navigation, the map should support smooth, reliable transitions; if long-range jumps dominated the code indiscriminately, they could corrupt local decoding and destabilize behavior. Sparse shortcut-carrying subpopulations allow nonlocal associations to remain available without disrupting the local geometry of the map. Thus, the same representational architecture may support two complementary regimes: broad population activity can preserve a locally accurate manifold for continuous traversal, whereas selective engagement or readout of multi-field subpopulations can expose long-range links for nonlocal access. A natural question is whether this shortcut overlay is actually engaged during hippocampal population events.

### 6.3 Synchronous events traverse the shortcut scaffold

A small-world scaffold is only meaningful if the system can and does traverse it. Prior work has shown that hippocampal activity can express nonlocal representations both online and offline [3,57,58]. During behavior, prospective sweeps have been reported at decision points [59, 60] and goal-directed navigation, where theta sequences reflect current goals and rapidly cycle among possible futures [57, 61–63]. During immobility and sleep, sharp-wave ripple (SWR) events support diverse forms of sequential reactivation [31], including forward and reverse sequences [64–66], reactivation of spatially and temporally remote experiences [58, 67], construction of never-experienced trajectories [59], anticipatory sequences preceding novel exploration [68], and sequences linked to goal-directed planning [2, 69]. In our calcium imaging data, immobility-associated synchronous population events align with the small-world scaffold, indicating that the scaffold is accessible. Because calcium imaging does not directly measure ripple oscillations, we treat these events as population-level phenomena that may overlap with replay-associated dynamics under appropriate conditions, rather than equating them one-to-one with electrophysiologically defined SWRs. This approach is consistent with previous studies [32, 33, 70].

Recent work emphasizes that replay spans multiple dynamical phenotypes [71]: stationary replay, where decoded positions remain at a single location [72]; diffusive replay, where decoded positions spread gradually through nearby states like Brownian motion [73]; and super-diffusive replay, where abrupt transitions carry decoded positions farther than random diffusion would predict [34,35]. Super-diffusive trajectories with intermittent long jumps are reminiscent of heavy-tailed step statistics often described as Lévy walks in animal search behavior [74, 75]. Lévy-walk-like dynamics have also been reported in neural population activity [76], and in our framework could arise naturally from mostly local traversal punctuated by rare wormhole hops. Consistent with this picture, we observe synchronous events that span the full range from smooth local progressions to abrupt jumps across distant locations and days, with the latter resembling super-diffusive dynamics expressed at the coarser temporal resolution of calcium imaging. More broadly, our framework offers a way to interpret and categorize heterogeneous ripple-associated population events by relating their decoded trajectories to an explicit representational scaffold, potentially helping to identify additional, previously uncharacterized event phenotypes.

How the brain selects between local traversal and shortcut access remains open. One possibility is that brain state changes which components of the hippocampal population participate in the active ensemble. Neuromodulatory signals [77], shifts in excitation–inhibition balance [78, 79], or changes in oscillatory regime [31] could bias activity toward locally continuous representations or toward subpopulations carrying nonlocal associations. Recent computational work further proposes that entorhinal input can tune hippocampal sequence generation between locally diffusive dynamics and occasional long jumps [80]. Plasticity may also shape the shortcut overlay itself; for example, behavioral time scale synaptic plasticity (BTSP) can assign credit over seconds and could help establish or reinforce sparse long-range links [81]. Finally, downstream circuits may implement flexible readout, either integrating hippocampal activity broadly to follow the smooth local geometry of the manifold, or selectively weighting subpopulations carrying multi-location associations to access long-range structure. Evidence that distinct hippocampal reactivation types are accompanied by distinct prefrontal modulation patterns supports the plausibility of such flexible readout [82].

### 6.4 Implications for world models and artificial intelligence

The organizational principles we identify may inform the design of artificial world models and reasoning systems to better plan, reason, and generalize. Large language models have demonstrated remarkable capabilities, but often require extensive test-time computation (chain-of-thought reasoning, tree search, or iterative refinement) [83–88] to solve complex problems. World models that learn latent dynamics for planning [89–92] face similar challenges: rolling out trajectories through a learned latent space can be computationally expensive, especially for long-horizon, complex tasks. Our results suggest a concrete architectural hypothesis: efficient search emerges when representations have reliable local neighborhoods and sparse nonlocal bridges that reduce the number of steps to reach distant states. For retrieval-augmented systems that perform multi-hop access over large knowledge stores [93, 94], it motivates regularizing learned graphs toward high local clustering with shorter global paths.

The clone-structured causal graph (CSCG) [95–99] provides a compelling framework in this context. CSCG addresses perceptual aliasing by learning a graph where “clones” of sensory states represent the same observation in different temporal contexts, explaining phenomena like splitter cells and context-dependent place coding. Recent work has shown that CSCG uniquely reproduces not only the final orthogonalized state representations observed in mouse hippocampus, but also the trajectory of learning, unlike sequence models such as LSTMs or transformers [13]. We note that our work follows the cognitive map’s geometry as it develops over learning, characterizing its reorganization along specific axes—rotation, drift, and the emergence of long-range links. How the underlying state structure itself is acquired is addressed by CSCG, which models the formation of an orthogonalized, aliasing-resolved state graph from experience [13, 95]. Schema-based extensions suggest how such graphs can be reused under higher-level abstract structure [99, 100]. Our results suggest targeted extensions for planning: add a separate sparse set of long-range “portal” edges between clones that reliably co-occur across distant contexts (analogous to coordinated multi-field structure) to impose a small-world prior, and gate portal influence based on task demands or uncertainty [97, 98].

The generalized condensed representation offers a further architectural parallel. In transformers [101], each token embedding compresses information from a context window into a current representation, enabling downstream computation without explicit retrieval of every prior token. Similarly, helical dynamics condense information about each state’s local temporal neighborhood. This can be viewed as a geometric “context window”: each point on the manifold implicitly represents not just a single state but a neighborhood of temporally related states, packaged in a form that downstream circuits can use directly. This structured and dynamic compression may enable efficient readout and computation in downstream areas without requiring explicit replay or retrieval of the full sequence.

### 6.5 Outlook

A key question is how shortcut structure is controlled during behavior. Efficient planning requires controlling when to exploit shortcuts versus smooth local traversal. Tasks can be designed to dissociate local accuracy from global access: for example, requiring choices among multiple remote candidates that are not locally distinguishable. Simultaneous recordings of hippocampus, entorhinal cortex, and prefrontal cortex can test whether shortcut engagement depends on available abstraction scale and whether prefrontal signals bias local *versus* global traversal. Causal perturbations can target population-level structure, including closed-loop disruption of immobility-associated events [79, 102] and manipulations that selectively weaken coordinated multi-field structure while preserving local decoding fidelity.

Small-world structure can also approximate a hierarchical graph that enables coarse-to-fine routing. Work on decentralized navigation suggests that hierarchy efficiency depends on distance-dependent rules for shortcut lengths distribution (often close to a power law) making simple local, greedy steps reliably find short routes [103], which may be followed in the cognitive map. The hippocampal-entorhinal system offers natural multiscale building blocks for such an organization [104]: grid cells are arranged into modules with distinct spatial periods [105, 106] and representational scale varies systematically along the hippocampal long axis, including gradients in place-field size [107–109].

More broadly, treating representational topology as a shared design variable offers a productive path forward for the continuing exchange between neuroscience and AI. In artificial systems, shortcut density can be manipulated directly and evaluated for reasoning and robustness. In neuroscience, a rapidly maturing suite of manifold and latent-variable methods [20, 47, 110–113] reveal increasingly articulate population-level structure of neural codes across brain regions whose topology reflects computational demands [16, 114–116]. Testing topological predictions in neural circuits and translating the resulting principles into world-model architectures may yield insights in both directions.

## 7 Methods

### 7.1 Experimental dataset and preprocessing

All analyses were performed on a previously published, publicly available longitudinal two-photon calcium imaging dataset from hippocampal CA1 collected during learning of a memory-guided virtual navigation task [13]. In that study, male and female Thy1-GCaMP6f mice were head-fixed in a virtual-reality setup and trained on the 2ACDC task, in which an indicator cue early in a 230 cm corridor signaled whether reward would be delivered at a near or far reward cue later in the trial. Imaging data are available via Figshare (https://doi.org/10.25378/janelia.27273552), and an interactive visualization is provided at cognitivemap.janelia.org.

We used the neural and behavioral variables provided with the release, including session-aligned behavioral position and trial type, as well as the processed neural activity traces for CA1 neurons tracked across sessions as described in [13]. Briefly, in the original work neurons were recorded with a two-photon random access mesoscope at 10 Hz across multiple adjacent imaging regions, and cells were registered across days using image registration and clustering procedures, with fluorescence traces extracted using Suite2p [117]. All analyses used deconvolved calcium traces extracted by Suite2p. Unless otherwise noted, our analyses focus on the core near- and far-reward trial types from the standard 2ACDC task and use the author-provided preprocessing and cell tracking without re-segmentation.

### 7.2 Γ-Autoencoder Architecture

Like many other techniques, Γ-AE [20] constructs a low-dimensional nonlinear manifold of the data. Unlike other techniques, however, Γ-AE selectively penalizes all distortions involved in the manifold geometry. We provide a discussion about Γ-AE in comparison to other techniques below.

An ambitious goal is to use an autoencoder to construct a low-dimensional and interpretable model of the mouse’s cognitive map that elucidates the most salient variables and accurately represents the quantitative relationships between them. A common failure mode of many deep neural networks is that they are universal function approximators that form latent spaces whose projections into the space of neuron activity are highly distorted [118, 119]. Therefore, in the models they produce, distances and angles in the latent space are not preserved on the manifold in the space of neuron activity, making it difficult to formalize quantitative relationships between the discovered variables (Fig. 1d). In addition, small changes along a latent space coordinate can lead to wildly fluctuating neural firing patterns, which makes it difficult to understand what moving along a latent space coordinate means in terms of changes to the neuron firing patterns. Our geometry-preserving autoencoder, Γ-AE, corrects these distortions by adding additional terms in the loss function that selectively penalizes excess manifold nonlinearities (see below). This procedure produces a model manifold with enough nonlinearity to quantitatively represent the data while locally preserving distances, angles, and directions when mapping latent coordinates to the manifold in data space. Preservation of distances ensures that small (large) changes in the latent space correspond to small (large) changes in the pattern of neuron activity. Preservation of angles ensures that directions that are orthogonal in the latent space correspond to changes in neural firing patterns that are maximally independent. Finally, preservation of direction ensures that moving along a particular direction in the latent space leads to regular and consistent changes of neuron activity. By including these costs in the autoencoder, we construct a model of the mouse’s cognitive map as a low-dimensional manifold that preserves the quantitative structure in the population activity, allowing us to study how the map is organized to support efficient search (Fig. 1h).

We achieve this manifold model by completely and orthogonally decomposing all manifold non-linearity into two curvatures: the extrinsic curvature that measures how quickly the direction of the collective variables changes along the manifold coordinates, and the parameter effects curvature that measures distortions in the distances and angles between the latent space and the data space. Hence, Γ-AE enables us to construct nonlinear manifolds that are interpretable—directions in the latent space correspond to consistent directions in data space—and quantitative—distances and angles in the latent space reflect those of the manifold in data space—even in regions with sparse or no data. As a result, Γ-AE builds a geometrically consistent manifold not just of the data, but of the entire nonlinear sub-space that the data inhabit.

To build our model manifold of hippocampal CA1 neuron activity, we use a deep neural network called an autoencoder comprised of two feedforward multilayer perceptrons: the *encoder*, ***f***_***θ***_: ℝ^*N*^ → ℝ^*k*^, and the *decoder*, ***h***_***ϕ***_: ℝ^*k*^ → ℝ^*N*^. The encoder architecture has four layers, where the layer dimensions are 200, 100, 50, 3, and the decoder architecture has four layers, where the layer dimensions are 3, 50, 100, 200. Hence, the function for the encoder follows

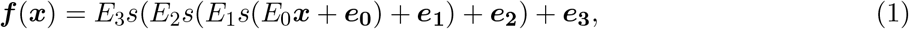

where the trainable parameters, ***θ***, are linear matrices *E*_0_, *E*_1_, *E*_2_ and bias terms ***e***_**0**_, ***e***_**1**_, ***e***_**2**_, and ***e***_**3**_. The activation function *s* is softplus. The function for the decoder follows:

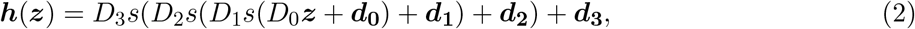

where the trainable parameters, ***ϕ***, are linear matrices *D*_0_, *D*_1_, *D*_2_, and *D*_3_ and bias terms ***d***_**0**_, ***d***_**1**_, ***d***_**2**_, and ***d***_**3**_. Passing an input vector of neuron activity ***x*** produces the output of the autoencoder 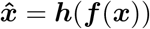.

### 7.3 Manifold Geometry

Because deep neural networks are universal function approximators, the manifold formed by the decoder can, in principle, distort to pass through the data in any arbitrary order. According to [20], we orthogonally decompose and regularize all manifold nonlinearities to selectively choose which nonlinearities to allow. This decomposition is comprised of two components.

First, to correct for distortions in the distances and angles between the latent-space coordinates and the manifold, we regularize the *parameter-effects curvature*:

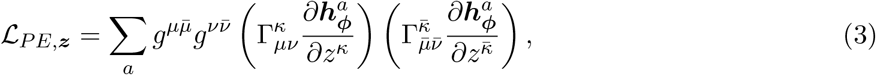

where *g*^*µν*^ is the inverse of the metric tensor, 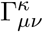 are the Christoffel symbols [120], and Eq. 3 is written in Einstein summation notation (repeated Greek indices are summed). The parameter-effects curvature given by Eq. 3 is the norm of the component of the second derivative of the manifold that lies tangent to the manifold at that point. Intuitively, Eq. 3 measures by how much a regular grid of points in latent space is distorted on the manifold by the decoder in data space.

Second, to ensure that the manifold coordinates are interpretable such that consistent directions in the latent-space correspond to similar changes in the patterns of neuron activity, we enforce that straight lines in latent space correspond roughly to straight lines in data space by also regularizing the *extrinsic curvature* at each point on the manifold, which we derive as

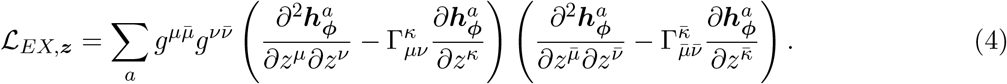

The extrinsic curvature is the norm of the component of the second derivative of the manifold that lies normal to the manifold at that point. Intuitively, Eq. 4 measures by how much the manifold tangent bends into new directions as we traverse the manifold.

Additionally, even though Eq. 3 quantifies nonlinearities on the manifold tangents, there is still an arbitrariness to the absolute global scale between the latent-space coordinates and manifold tangents (e.g. how far one unit distance in the latent space is on the manifold). To fix an absolute scale, we include an additional loss on the scale of the manifold to be approximately uniform such that

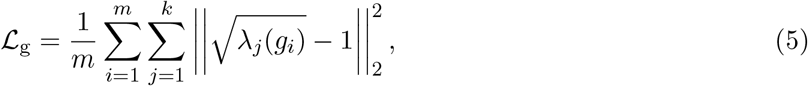

where *g*_*i*_ is the metric tensor at sampled points *i* in the latent space. Additionally, while Eq. 3 and Eq. 4 define the distortions of the manifold geometry through the decoder (Eq. 2), we additionally regularize distortions for the encoder ℒ_*E*_ that maps points from the neuron space onto the latent space.

Finally, the network also minimizes the standard autoencoder reconstruction loss,

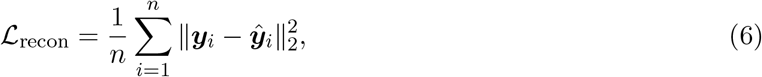

which is the mean squared error of reconstructing the original data from the manifold. These losses combine to yield the final loss

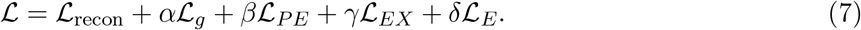

In practice, we want our manifold to preserve distances and angles as well as possible, so we set *α* = *β* = *δ* = 100 to be very large. Hence, the only nonlinearity we allow meaningful quantities of is the extrinsic curvature *γ* = 0.5, which allows the manifold to bend gently along with the natural curvature of the data.

### 7.4 Constructing the surfaces of experience

Once Γ-AE is trained, we use the data points projected in the latent space to construct our surfaces of experience. To build each surface for each task condition, we average the latent space points corresponding to every 3 cm of track and every 20 trials for that task condition. Once the data are binned in a grid in the latent space, we smooth the surfaces using taubin smoothing [121] that avoids the artifact of shrinking the manifold experienced by Gaussian smoothing. We used taubin smoothing parameters *λ* = 0.2 and *µ* = −1.01*λ*, and iterated until the algorithm converged.

### 7.5 Computing the generalized state fields

In this work, we define a GSF for a neuron as localized regions of the latent space that have high decoded activity for that neuron. To compute GSFs, we first densely and uniformly sampled the latent space using a regularly spaced grid of points, *X*, such that the grid extends slightly beyond the extent of the observed data embedded in the latent space. Then, we constructed a mask containing all grid points within the convex hull of the observed data. The points inside of the convex hull were decoded to patterns of neural activity, whereas the corresponding activity of points outside of the convex hull were set to 0 for all neurons. Then, each neuron’s activity on the grid was smoothed using a Gaussian convolution with a kernel standard deviation roughly 1/10th of the standard deviation of the observed embedded data. For clarity, we will call this Gaussian convolved activity for each neuron *n* from the decoded latent space grid the *latent activity map*, denoted *L*_*n*_(***x***), ***x*** ∈ *X*. For the visualization of the GSF isosurfaces, we threshold each isosurface to show activity above 98.5% of the activity for that neuron on the latent space grid. This threshold was used purely for visualization, and not for any quantitative analysis.

### 7.6 Computing statistically significant GSF peaks

To go beyond a thresholded visualization and define statistically significant peaks of the GSFs for further analysis, we begin with the latent activity map for one neuron, and identify candidate peaks as local maxima. Then for each peak, we use a merge-tree algorithm to compute its *prominence* as the difference between that neuron’s activity at that peak and the highest saddle point on a path connecting to any higher peak. Said another way, the prominence for any peak is the minimum decrease in that neuron’s decoded activity to reach a higher point. The maximum peak is given a prominence of infinity. Given each peak *i* of neuron *n*, the decoded activity of that peak, *a*_*i*_, and the prominence of that peak, *b*_*i*_, we consider a candidate GSF peak to be a true significant GSF peak if both its activity relative to the minimum decoded activity in the latent activity map, and its prominence, satisfy

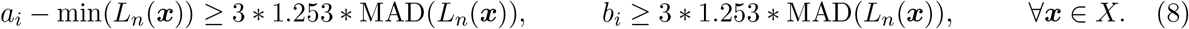

Here, MAD is the mean absolute deviation, and the factor 1.253 is the scale between the MAD and the standard deviation for normally distributed data. We define the set of all statistically significant peaks that pass these criteria as 𝒫 = {*p*} with coordinates ***x***_*p*_ and the corresponding neuron that produced that peak *n*_*p*_.

### 7.7 Computing statistically significant GSF co-peaks

Once we have the set of statistically significant GSF peaks, we turn to statistically testing for significant co-peaks. To do this, we take the set of all statistically significant GSF peaks across all neurons, 𝒫, choose one peak at a time as the query peak, *p*_*i*_ ∈ 𝒫, find the *k* = 200 nearest neighbor peaks, 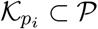, identify which neurons gave rise to those neighboring peaks, 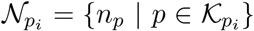, and find all peaks produced by those neurons 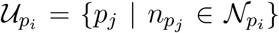. The candidate co-peaks, then, are the subset of peaks produced by those neighboring neurons excluding the neighboring peaks, 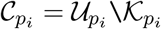, and 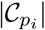 is the number of candidate co-peaks.

To test whether the co-peaks are significantly clustered relative to chance, we perform a shuffle test and look at the distance from each co-peak to its *m* = 30 nearest neighbor. Specifically, for each co-peak 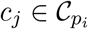, we compute the distance to the *m*-th nearest co-peak, *d*_*j*_. Then, we select a random draw of 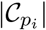 peaks from all significant peaks excluding the neighboring peaks, 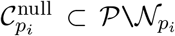, and compute the distance from each true co-peak *c*_*j*_ to the *m*-th nearest null co-peak, 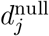. Finally, we define a true co-peak *c*_*j*_ to be statistically significantly clustered to other true co-peaks than by chance if *d*_*j*_ is smaller than 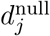 for all of 5000 random draws of 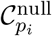, the set of which we define as 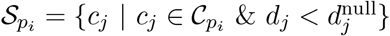.

### 7.8 Defining small-worldness using latent-space co-peaks

The procedure for defining statistically significant GSF co-peaks associates each query point *p*_*i*_ ∈ 𝒫 with the set of significantly clustered co-peaks 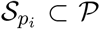. To define small-worldness in this space, the fine-grained approach would be to consider each peak as a node and define an edge as existing from node *p*_*i*_ to *p*_*j*_ if the significant co-peaks 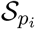 overlapped with the neighborhood 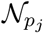. This procedure would produce a graph of |𝒫| nodes. To produce a coarser graph, we take a subset of this graph where the nodes in that subset are chosen sequentially via a greedy set cover such that the neighborhoods of the chosen points maximally cover all of the peaks. It is this graph subset that is shown in the main text.

### 7.9 Defining rate maps in position and day

To identify place field peaks in the raw data across position for each day, we first build rate maps for each neuron’s activity. To build these rate maps, for each neuron *i*, we first binned all frames into bins of 5cm and 1 day, such that for each position bin *b*, we get the summed activity *a*_*i*_(*b*), and the number of frames in that bin *n*_*i*_(*b*). To avoid large fluctuations from bins with few frames, we compute the rate as the ratio of the convolved summed activity and frame count

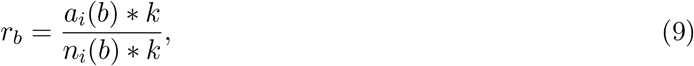

where ∗ denotes the convolution with the kernel *k*, which is a Gaussian with a standard deviation of 5cm. This rate map is constructed for each day and each task condition. In the context of the rate maps and subsequent analyses of significant peaks, we consider each task condition separately as a circular track, such that these convolutions are taken on two periodic tracks, producing a rate map for each task condition. We then concatenate these two rate maps together, and for any analysis involving the rate maps and subsequent peaks, refer to the bin index along this concatenated rate map as position.

### 7.10 Defining statistically significant rate map peaks

To define statistically significant peaks in the raw data across position for each neuron and each day, we construct the rate map for that neuron and day across position, and identify candidate peaks as local maxima in the rate map. To test whether these local maxima are statistically significant and robust, we perform a shuffle test. Specifically, we take all of the raw activity data as a function of the frame number, and for the frames of each trial, circularly shift the frame index by a random uniform integer between 1 and the number of frames for that trial. We perform this random shift for each of all trials. Then, we use this randomly shifted data to construct a null rate map 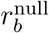 as if the frame indices were not permuted (i.e., *n*_*i*_(*b*) does not change, and only the data that goes into *a*_*i*_(*b*) come from the shifted null). For each null rate map, we compute the maximum value of 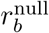, and repeat this null generation procedure 10,000 times. We define a rate map peak for each neuron and each day to be statistically significant if it is larger than 95% of the maximum null values, and also enforce a minimum peak distance of 2 bins.

### 7.11 Defining statistically significant co-firing peaks in the rate maps

To test for statistically significant co-firing peaks in the rate maps, we first construct a significant peak tensor *P* ∈ {0, 1}^#days × #position × #neurons^, where an index *P*_*ijk*_ is 1 if neuron *k* had a statistically significant peak on day *i* and position *j*, and 0 otherwise. Then, for a query day and position bin (*i, j*), we identify the neurons that fired in the immediate positional neighborhood of that query bin 𝒩_*ij*_ = {*k* | *P*_*itk*_ = 1, *t* ∈ {*j* −1, *j, j* +1}}, and computed how many of those neurons had significant peaks for each position and day bin, 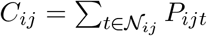, where we set the counts within the query bin and immediate positional neighborhood to 0. We refer to these counts as the empirically observed peak count.

To test whether an empirically observed peak count *C*_*ij*_ was statistically significantly greater than by chance, we performed Monte Carlo sampling from a multinomial null distribution with empirically observed probabilities. Specifically, for each day *i* and position *j*, we compute the number of significant peaks across all neurons as *D*_*ij*_ = ∑_*k*_ *P*_*ijk*_, and normalize those counts as the multinomial null probabilities,

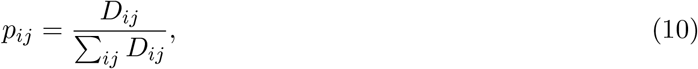

We then performed 10,000 draws from this multinomial distribution using *p*_*ij*_ as the multinomial probabilities, and ∑_*ij*_ *C*_*ij*_ as the total observed counts, and for each multinomial null draw computed the maximum standardized bin count, 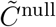. We considered an empirically observed standardized bin count 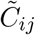 to be statistically significant if it was greater than 95% of the maximum null draws 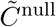. Hence, for each day and position bin, (*i, j*), this procedure defines a set of bins 𝒮_*ij*_ where neurons that fire in the positional neighborhood of (*i, j*) have statistically significantly more peaks than by chance. We convert these sets of significant co-firing bins into a matrix *S* ∈ {0, 1}^(#days∗#position) × (#days∗#position)^, where each row and each column corresponds to a day and position bin (i,j), and the entry *S*_*mn*_ = 1 if the query day and position bin (*i, j*) corresponding to index *m* has a significant bin in 𝒮_*ij*_ corresponding to index *n*.

### 7.12 Computing the small-world coefficient and generating the degree-preserving null model

To test whether our graphs comprising nodes of bins of position (5cm) and time (1 day) and edges from query bins to other bins with statistically significantly higher co-peak counts, we compute the small-world coefficient [37], which takes the ratio of the graph’s clustering coefficient relative to a null, and the graph’s shortest path length relative to a null. To construct the null, we used the Maslov-Sneppen algorithm [122] that performs degree-preserving edge swaps to generate 1000 randomized null graphs using ∼10 swaps per edge. From our empirically measured graph and the corresponding nulls, we computed the small-world coefficient by taking the fraction between the ratio of empirical *versus* null clustering coefficient, and the empirical *versus* null characteristic path length. We then computed the null distribution of small-world coefficient by computing the same ratio between each null re-wired graph’s clustering coefficient *versus* that of the null ensemble and the null re-wired graph’s characteristic path length *versus* that of the null ensemble.

### 7.13 Decoding and testing synchronized calcium events

Synchronized calcium events (SCEs) were identified from stationary periods during the active task rather than during pre-session or post-session rest. This criterion captures moments when the animal has voluntarily paused mid-task during goal-directed navigation—brief pauses thought to be associated with memory consolidation and planning. We refer to these as SCEs rather than sharp-wave ripples (SWRs), as calcium imaging does not directly measure the underlying oscillatory dynamics, though previous work has established that such population events can co-occur with electrophysiologically measured SWRs. A frame was classified as an SCE if it satisfied the following criteria: (1) speed = 0 cm/s, (2) not during the inter-trial teleportation period, and (3) no licking at that frame or ±2 frames. Given the 10 Hz imaging rate, each frame corresponds to 100 ms. To ensure temporal separation between SCE frames and running frames used for manifold construction, SCE frames were required to be padded by at least two consecutive stationary frames on either side, while manifold frames required padding by at least two consecutive high-speed frames (speed *>* 5 cm/s). This ensured a minimum separation of 500 ms between SCE and manifold frames, minimizing signal bleed-through.

To decode the synchronized calcium events (SCEs), we first associated an averaged pattern of neuron activity with each position *j* (5cm) and day *i* by averaging the activity for the frames where the animal was at that particular position and day. This procedure gives us an average vector of activity ***a***_*ij*_, which we then normalize to yield ***b***_*ij*_ = ***a***_*ij*_*/*|***a***_*ij*_|. This procedure of defining bins is distinct from how we constructed the rate maps because the goal was not to identify robust peaks along the continuum of position, but rather to have an accurate and unbiased measurement of each position and day bin for decoding.

To decode the activity of each SCE *m*, we take the normalized vector of its pattern of activity, ***s***_*m*_, and reconstruct it using non-negative least squares with a ridge penalty

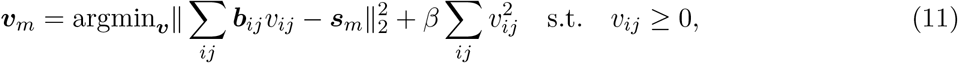

where we choose *β* = 1. This procedure gives us the reconstruction coefficients ***v***_*m*_ whose non-negative coefficient *v*_*ij*_ is the amount of weight given to the position and day bin ***b***_*ij*_ in the reconstruction.

To test whether the weights are statistically significantly large, we perform a shuffle test by randomly permuting the elements of ***s***_*m*_, and recomputing the coefficients 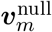 using the same procedure in Eq. 11 for 2000 permutations. For each shuffle, we record the maximum decoded coefficient. For SCE *m*, we call an empirically decoded coefficient statistically significant if it is larger than 95% of the distribution of 2000 maximum decoded coefficients from the null shuffles, giving us a thresholded vector ***t***_*m*_.

### 7.14 Testing overlap between SCE decoded maps and the small-world graph

To test whether the empirically decoded coefficients for the SCEs leverage the small-world graph more than by chance, we take the vector of significantly decoded coefficients for each SCE, ***t***_*m*_, and the matrix containing statistically significant co-firing peaks in the small-world graph *S*, and define traversing one step down the graph as

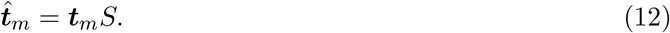

The alignment score, then, is the correlation between ***t***_*m*_ and 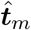.

To test whether this alignment is statistically significant, we construct a shuffle null where we take the decoded coefficients ***t***_*m*_ which contains coefficients for each day and position bin, and circularly shift the position index by a random integer between 1 and the number of positions - 1, and recomputed the overlap score. This circular shift ensures we preserve any spatial correlations so that our null overlap score is not artificially low due to breaking up spatial correlation. We perform this null shift 1000 times, and find that our empirically measured overlap score is larger than the entire null distribution by a wide margin.

## Data availability

All data used in this study are publicly available as part of the dataset released in [13] on Figshare (https://doi.org/10.25378/janelia.27273552). An interactive visualization of the dataset is available at cognitivemap.janelia.org [13].

## Code availability

Code used for analysis and figure generation will be released upon peer-reviewed acceptance.

## Acknowledgements

We thank Antonio Fernandez-Ruiz, Ian Ellwood, Chongxi Lai, Albert Lee, Melody X. Lim, Brett Mensh, and Nelson Spruston for valuable discussions and feedback on the manuscript. J.Z.K. was supported by postdoctoral fellowships from Bethe/KIC/Wilkins, Mong Neurotech, and the Eric and Wendy Schmidt AI in Science program of Schmidt Sciences, LLC. I.C. and J.Z.K. were supported in part by the NIH National Institute of Arthritis and Musculoskeletal and Skin Diseases (5R21AR083064-02) and the Air Force Office of Scientific Research (FA9550-23-1-0722). J.P.S. and I.C. acknowledge support from the National Science Foundation under Grant No. DMR-2327094. W.S. was supported by Cornell University.

## 8 Supplementary Information

### 8.1 Latent representation codes for position, task belief, and trials

To test the correspondence between the cognitive map models and the virtual maze we plot the projection of neural activity onto a 3-dimensional latent space learned by Γ-AE for a representative mouse in Figure 5 (see Methods). To identify the salient variables for which the hippocampus codes, we plot the embedding points for all frames on day 8, colored by the different track markers. For clarity, we plot the near trials opaque in Fig. 5a and far trials opaque in Fig. 5b. We observe that locally, data that vary along one direction in the latent space correspond to the mouse’s position along the track. We find that the cognitive map model arranges the neural firing patterns along the track in a manner that correctly organizes the maze marker sequence and topology for both task conditions. Additionally, we observe that, at each point, the data corresponding to the two tracks separate along a second direction throughout learning for the region spanning the indicator zone through the R2 zone (see Fig. 7 for the embeddings of the final day for all animals).

**Figure 5:**
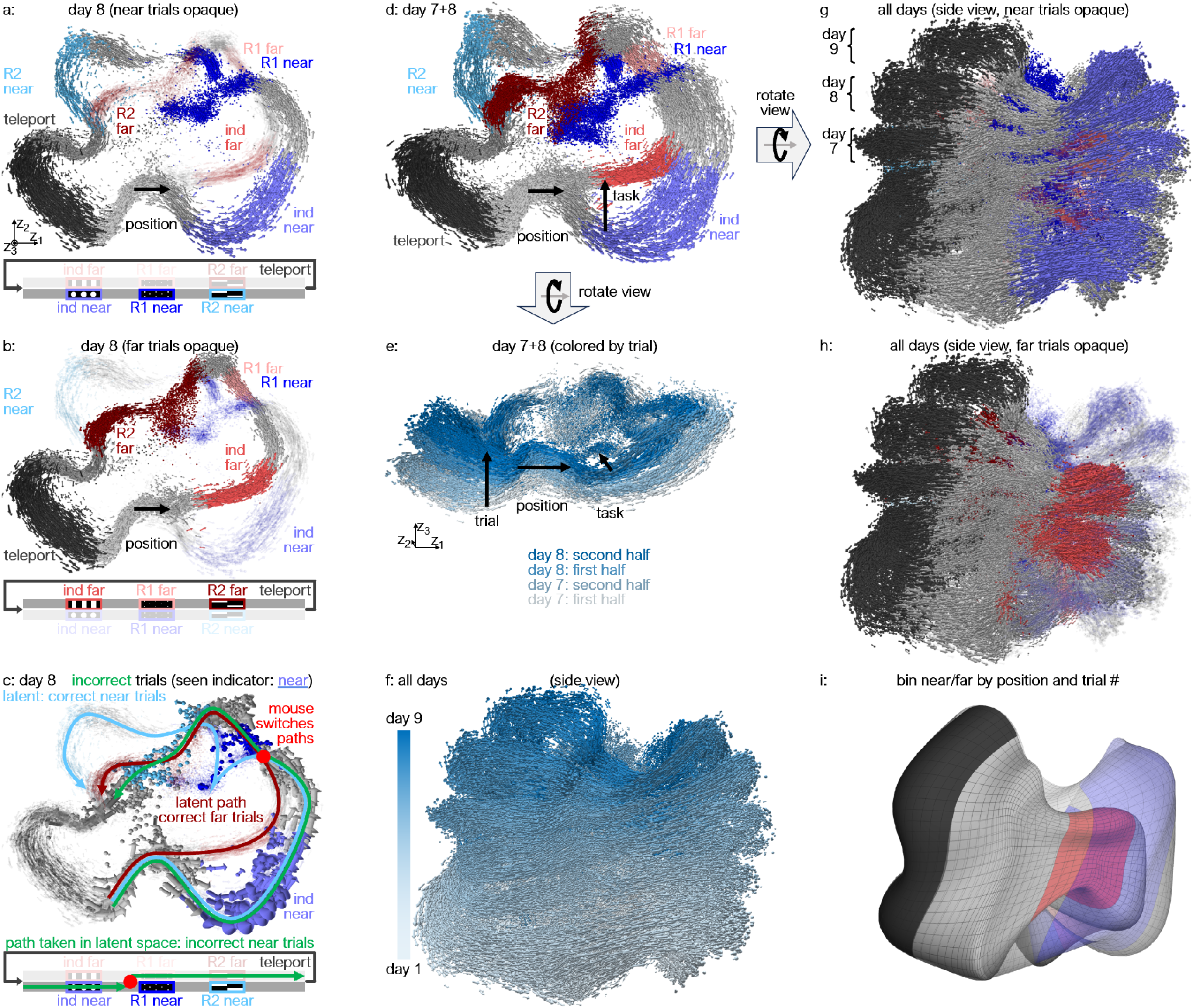
Low-dimensional geometry of neural representation along position, task, and trials. (**a**) Low-dimensional embedding of hippocampal activity and evolution from ∼5000 neurons in a 3-D latent space on day 8, colored by track marker, with data from the near trials opaque and the far trials transparent. (**b**) Same embedding with near trials transparent and far trials opaque. (**c**) Same embedding, with trials containing correct licking behavior transparent, and incorrect licking behavior during near trials opaque, showing most of the opaque points switching from the correct near trials to the correct far trials. (**d**) Embedding from days 7 and 8, colored by task marker showing near perfect overlap of track marker across days, and (**e**) rotated view showing a consistent direction of change in embedding coordinate within- and between-days, (**f**) which persists across all 9 days. (**g**) Embedding from all days colored by task marker with near trials opaque, and (**h**) far trials opaque, showing consistent alignment of task markers between all days. (**i**) Embedding data binned across position (3cm/bin) and trials (20 trials/bin) for near (transparent) and far (opaque) conditions.

**Figure 6:**
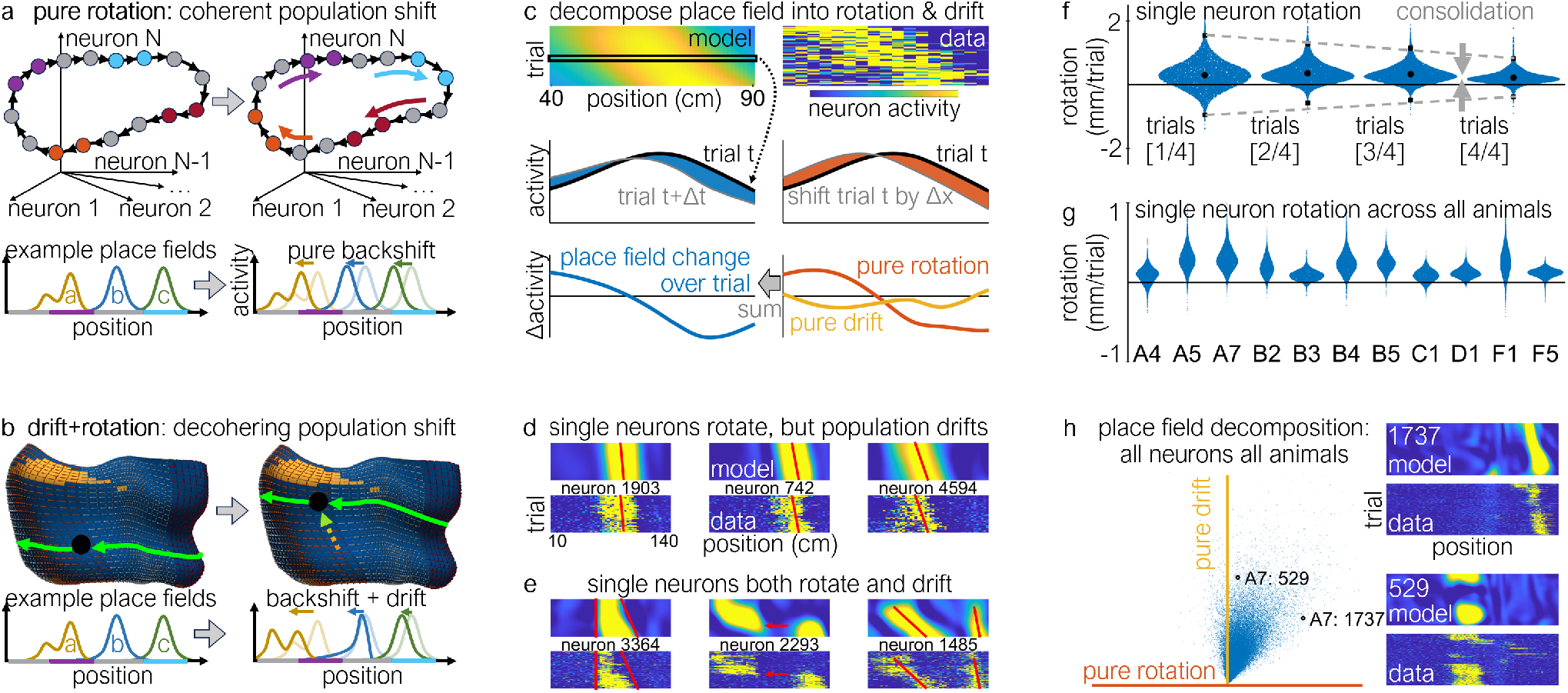
Individual place fields decompose consistently along rotation backshift and drift. (**a**) Schematic of a single trajectory in the space of neuron activity with points colored by track markers that shift as the representation undergoes pure rotation/backshift (top), and 3 corresponding examples of single-neuron place fields undergoing pure backshift (bottom). (**b**) Schematic of a single trajectory undergoing both rotation/backshift and drift on the binned manifold (top), and 3 corresponding examples of single-neuron place fields undergoing a combination of backshift and drift (bottom). (**c**) model and data of a single neuron’s activity over position and trial (top), the change in activity across trial (blue) and from a pure rotation/backshift (orange) from the model (middle), where the change in activity over trials can be decomposed as a weighted sum of the pure rotation and a pure drift (bottom). (**d**) Model prediction (top) and data (bottom) of activity for single neurons displaying mostly rotation, but at different rates, leading to population-level drift. (**e**) Single neurons displaying a complex combination of rotation and drift. (**f**) Consolidation of rotation/backshift distribution for single neurons over time. (**g**) Rotation/backshift across all neurons averaged across all trials consistently positive for all animals. (**h**) Phase diagram of average rotation and drift component for each neuron and animal over all trials with two example model and data activity maps.

**Figure 7:**
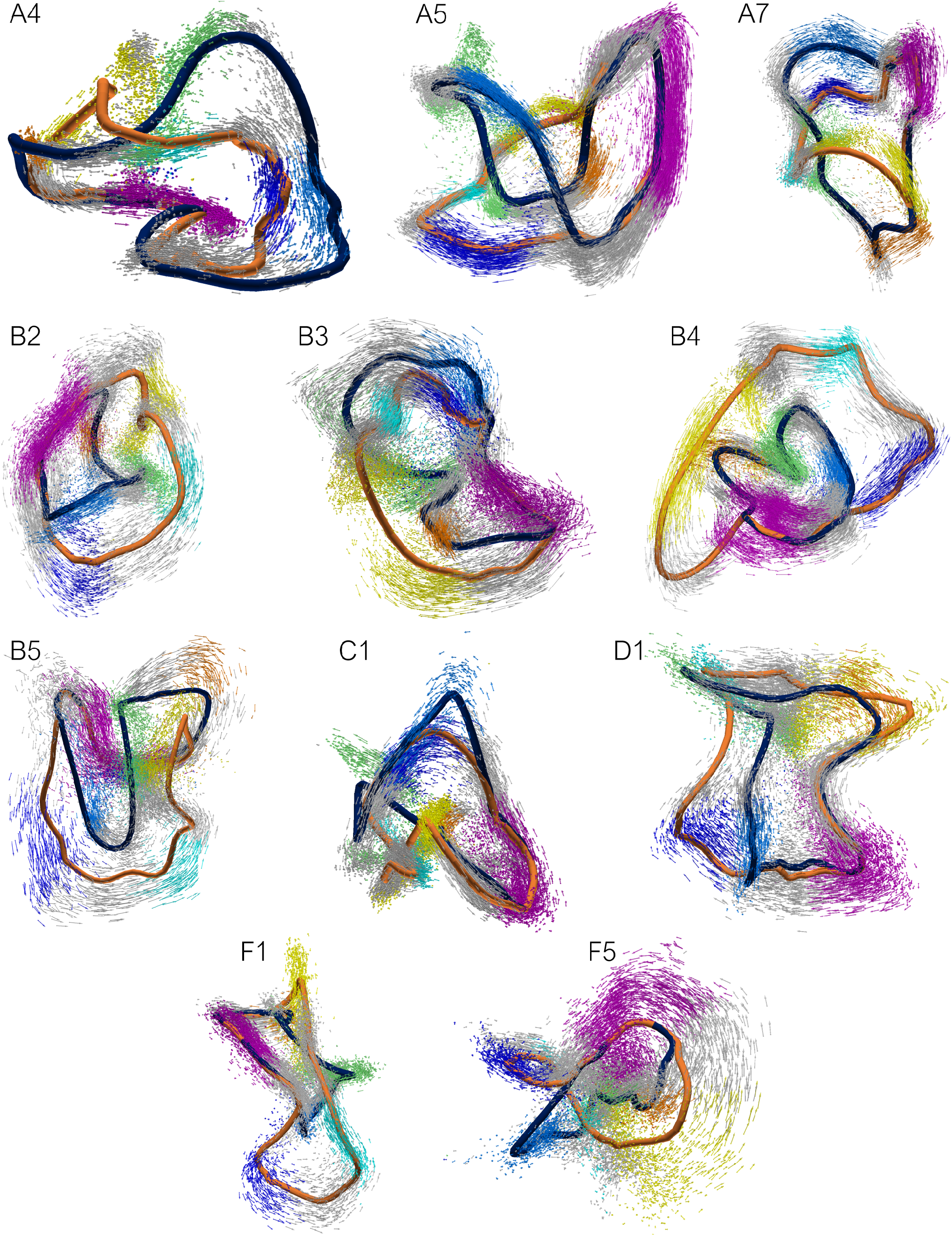
Embedding of final day’s representation for all animals. The embedding coordinates and vectors for the final day’s representation across all animals. Each vector is colored by the physical track marker of each mouse’s position on the linear maze, and the navy (orange) trace is the average trajectory for the near (far) trial on the final day.

**Figure 8:**
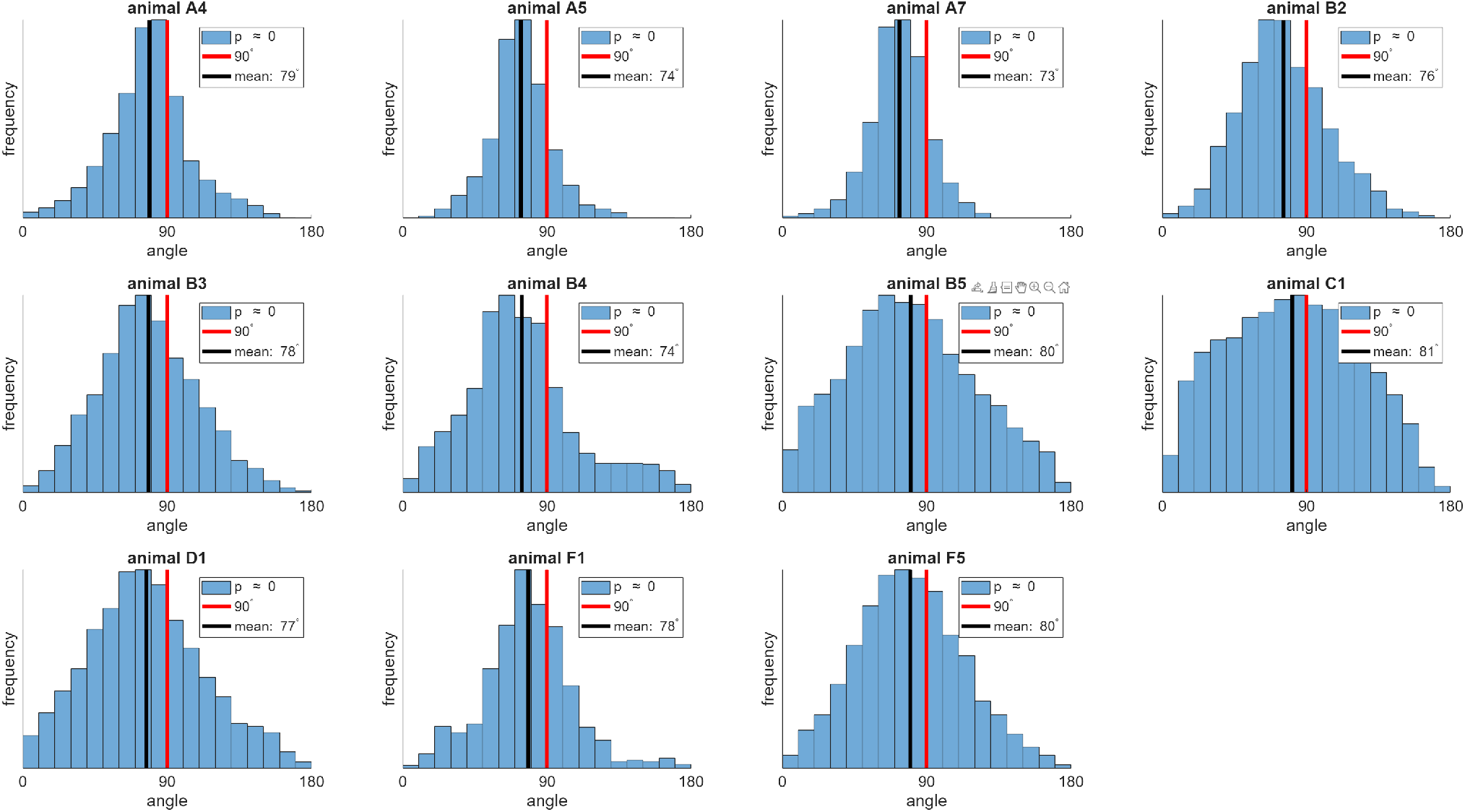
Distribution of angles between the position and trial vectors in the latent space for all animals, statistically tested using a 1-sided t-test against a null mean of 90°.

**Figure 9:**
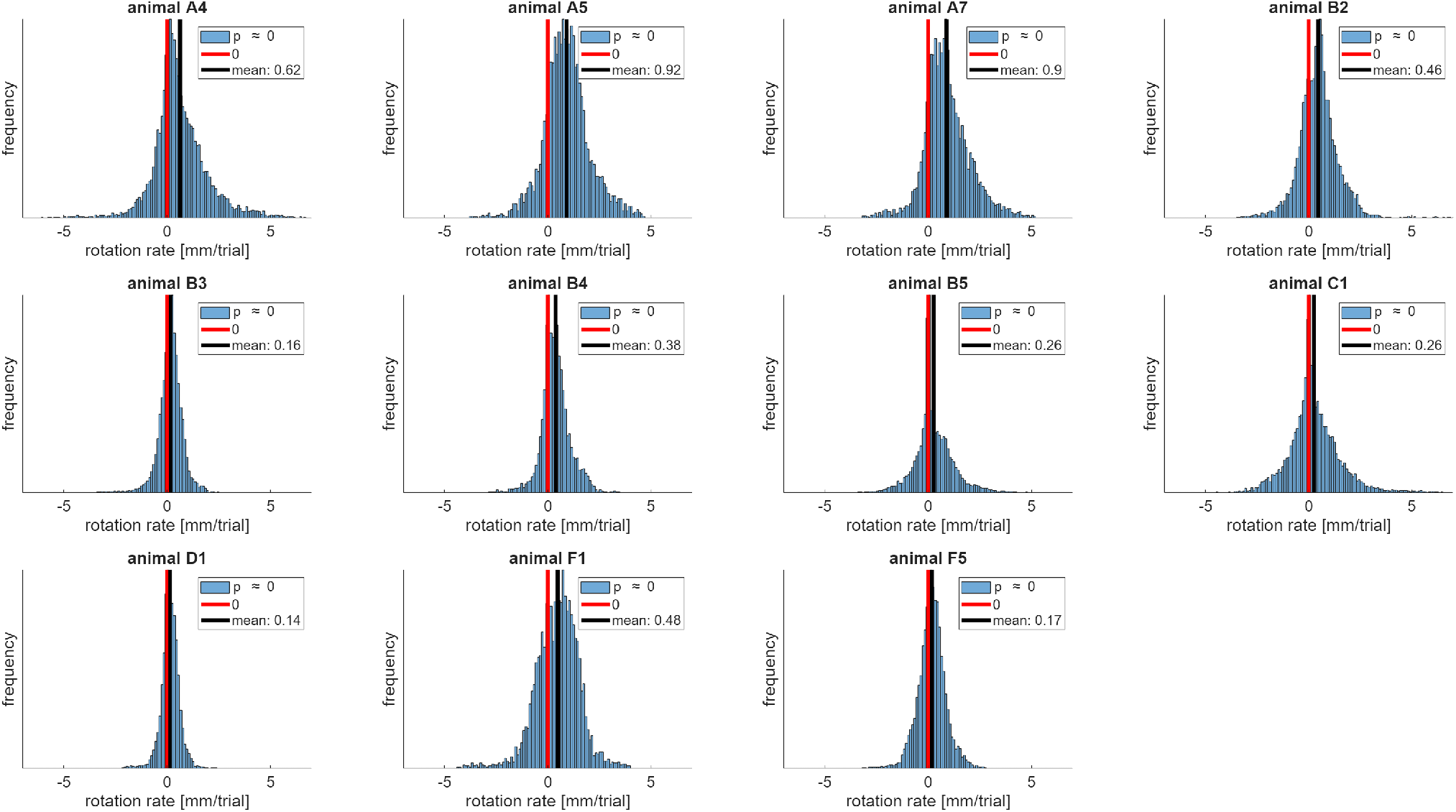
Distribution of the advancement in positional representation per trial induced by the rotation in the surfaces of experience in the latent space for all animals, statistically tested using a 1-sided t-test against a null mean of 0mm/trial.

**Figure 10:**
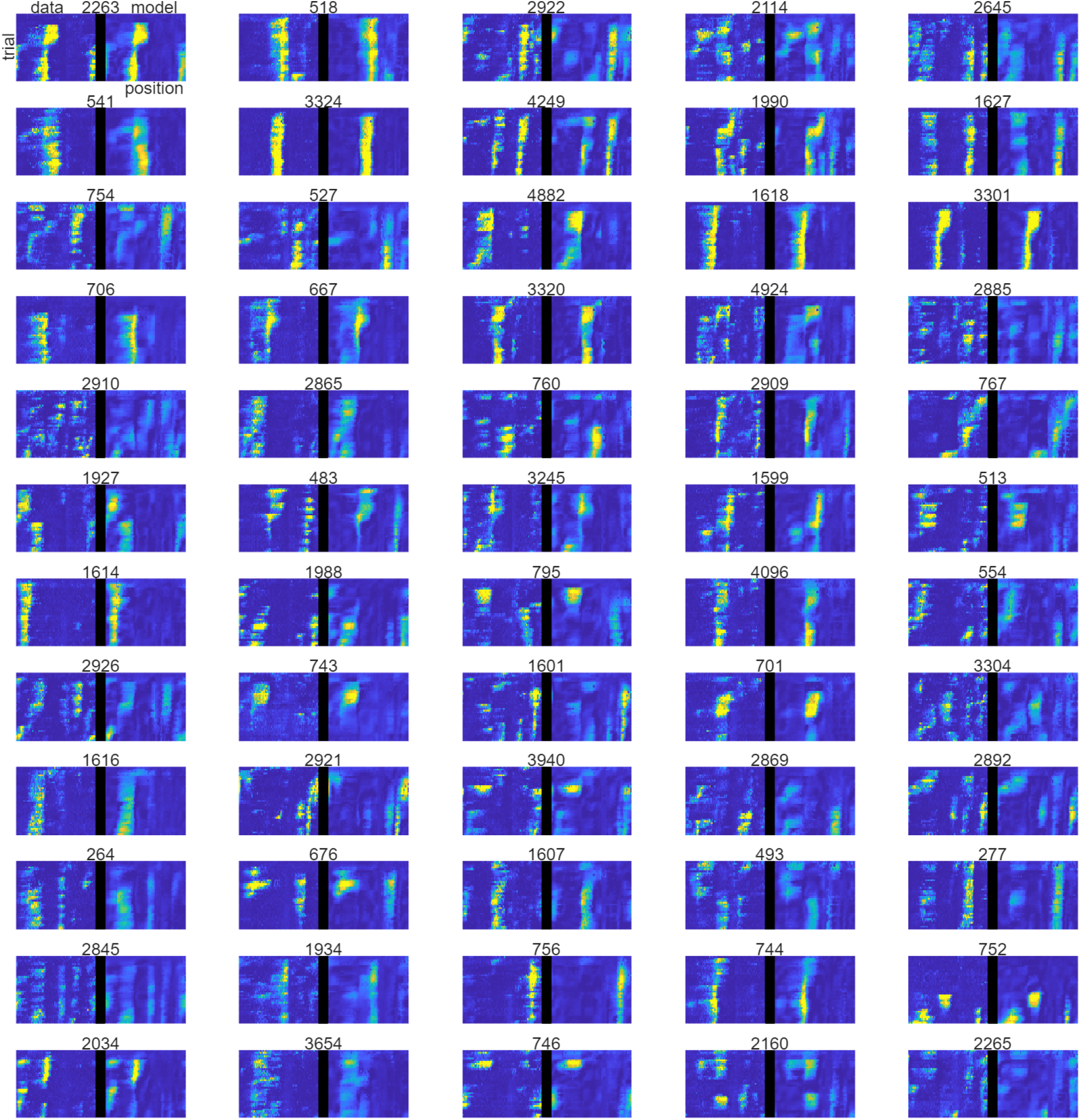
Data and model prediction of single neuron activity. The activity of the 60 neurons with highest mean activity from the data and the corresponding model prediction decoded from the 3D latent space. Each bin is 30mm by 20 trials across all trials and all days for animal A7.

**Figure 11:**
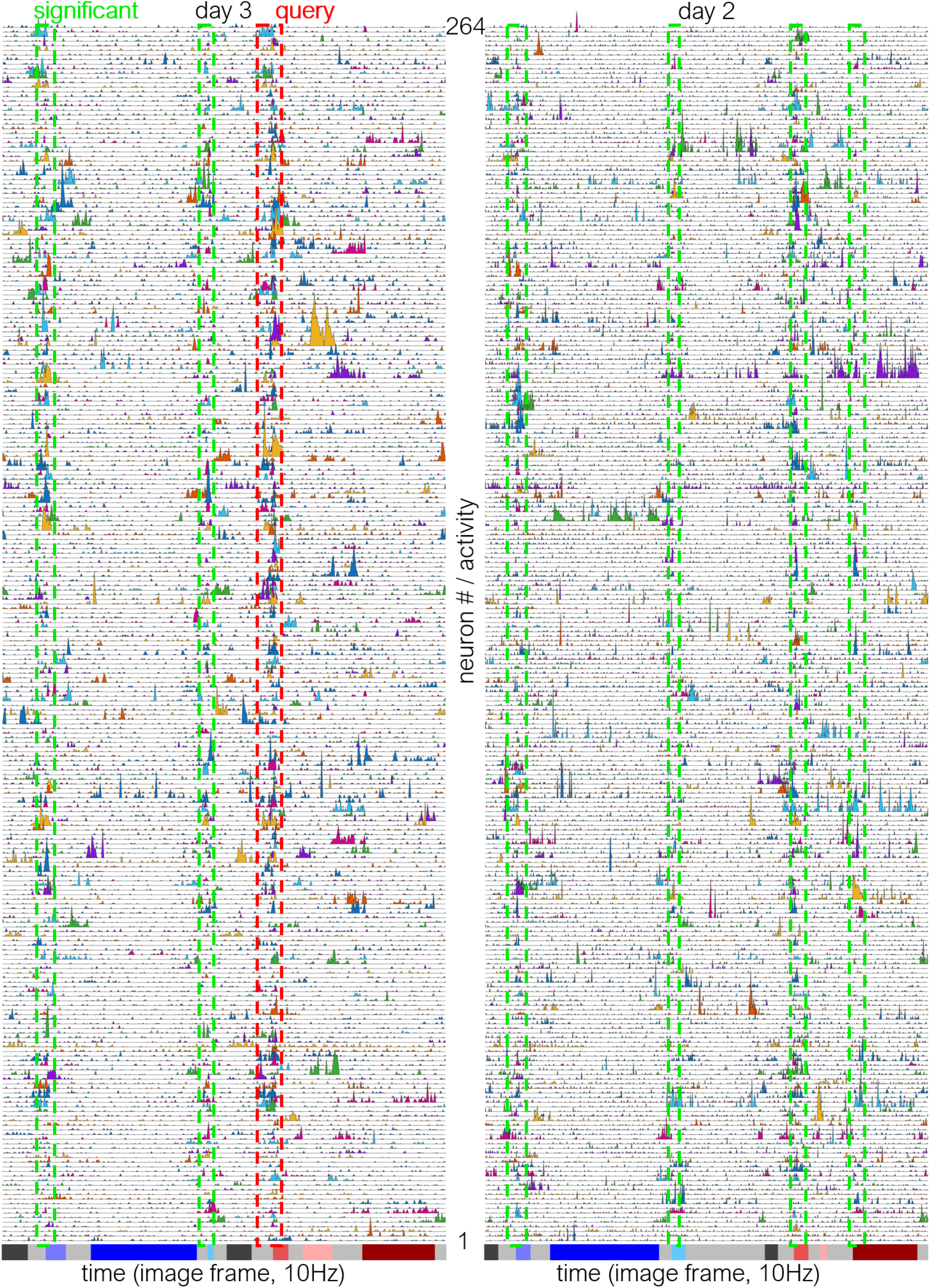
Raw unfiltered time traces of activity for 264 neurons in the main text Figure 4 for a consecutive near and far trial on day 3 (left), and another consecutive near and far trial on day 2 (right), with the query region roughly marked in the dashed red box, and significant co-firing regions roughly marked in the dashed green boxes.

**Figure 12:**
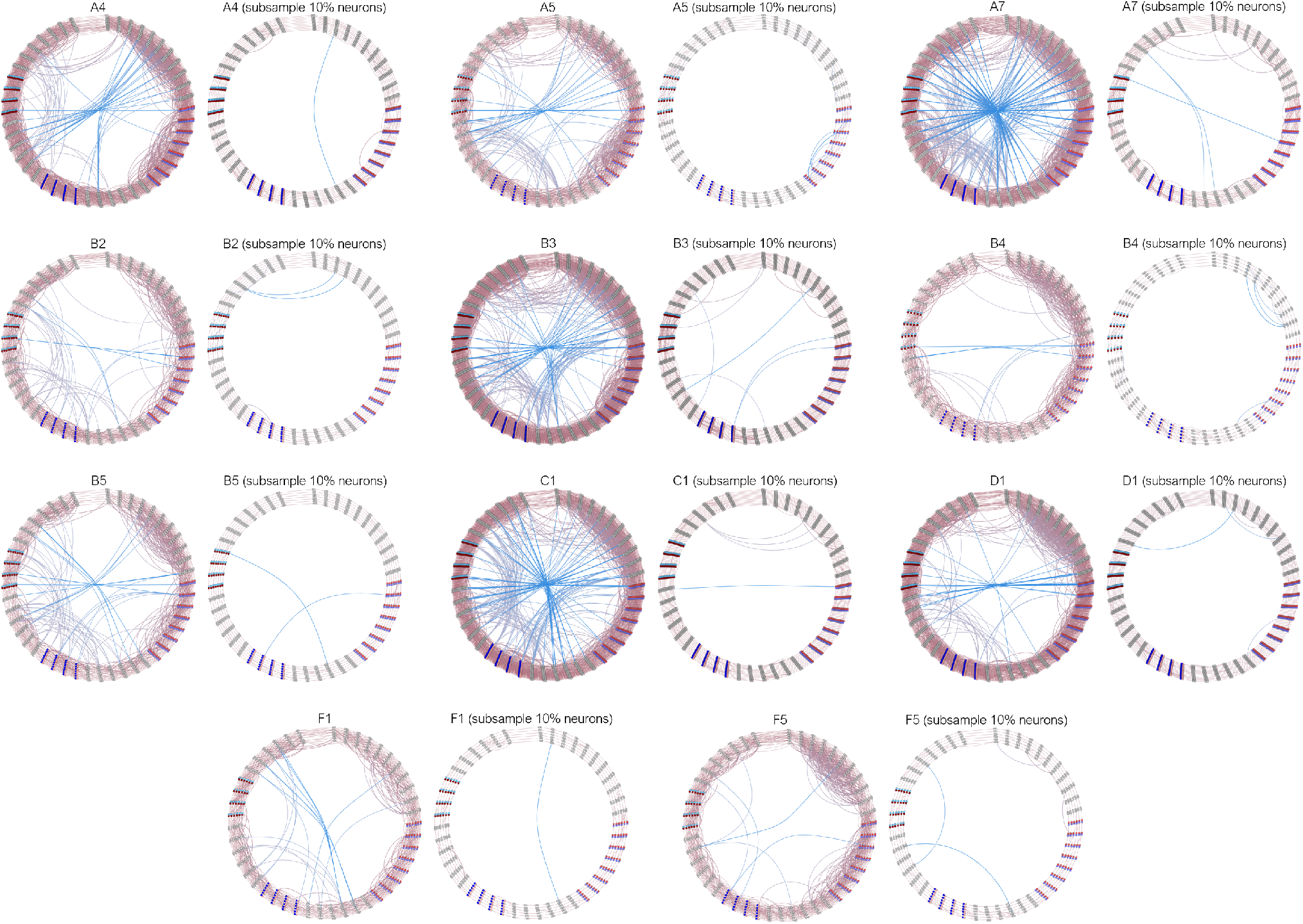
Small-world graphs of significant co-firing cells for all animals using all neurons, and using 10% of neurons.

**Figure 13:**
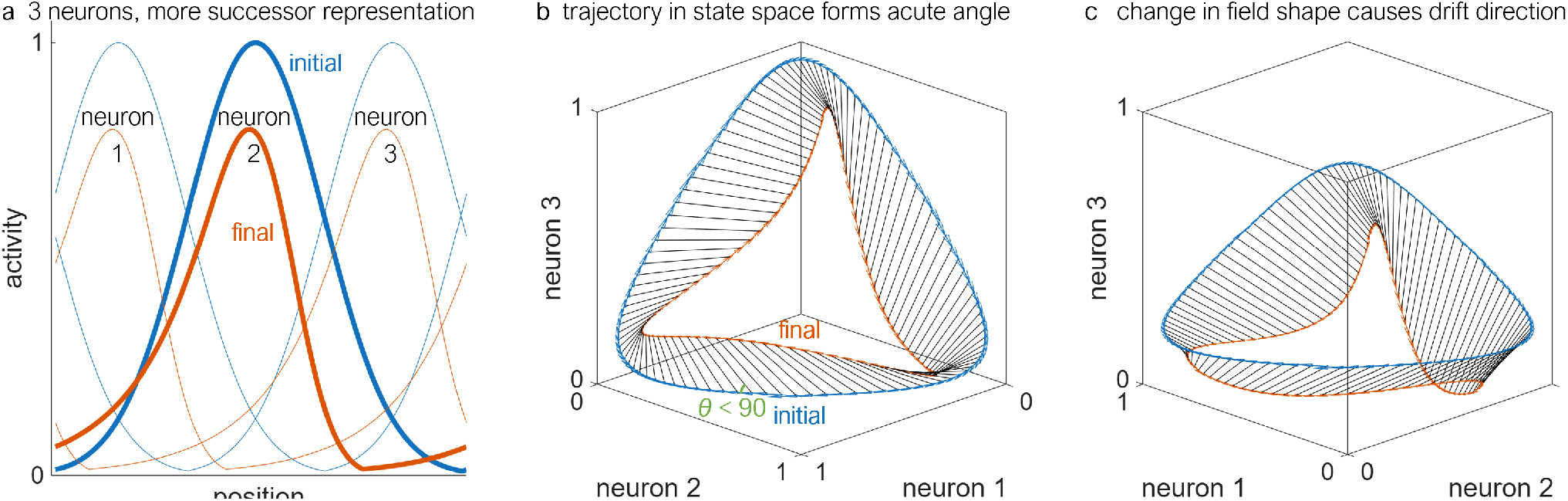
Geometry of single neuron place field under successor representation causes rotation and drift. (**a**) Example of three artificial neurons coding for a circular track with initially unbiased position tuning (blue) and eventual direction-biased tuning (orange). (**b**) Plot of both the initial and final trajectories in the 3-dimensional state-space of neuron activity, showing how the geometry of the track changes and creates an acute angle. (**c**) Same state-space plot of the two trajectories viewed at a different angle from the side, showing that the change in the individual field shape induces a drift in representation.

Interestingly, we also observe trials where the cognitive map model starts along a trajectory associated with one task condition, but switches to the trajectory associated with the second task condition (Fig. 5c). Such switches often occur during error trials (licking at the wrong reward zone) or when the mouse is being inattentive (running without licking). This path switching on the manifold accompanied by incorrect licking behavior provides evidence that the mouse is confused about the task condition it thinks it is in. Collectively, these results indicate that each pathway on the manifold corresponds to the task condition in which the mouse believes itself to be and that the separation between these tracks, a second local direction in the latent space coordinates, represents how much the mouse believes each portion of the track belongs to each task condition (task belief).

To identify an additional variable for which the hippocampus codes, we plot the embedding vectors for both days 7 and 8, and observe that all track markers and the track topology are strongly aligned between days (Fig. 5d), thereby largely preserving the first two dimensions of position and task condition. By rotating the viewing angle and coloring the points now by the first- and second-half of trials on each of days 7 and 8 (Fig. 5e), we observe that the data consistently vary along a third geometric axis as a function of trial number, which is preserved within and between days. By plotting all of the points from all days and coloring the points by the trial number, we observe a consistent and continuous gradient along this third geometric axis that persists across a whole week (Fig. 5f). By plotting all of the points and instead coloring them by the task markers, we additionally observe that the data from the near task conditions (Fig. 5g, opaque) and the far task conditions (Fig. 5h, opaque) remain separated along a direction of task belief through trials, which can be decoded to and associated with a unique pattern of neural activity. Collectively, these analyses demonstrate that a specific point on our model’s latent-space corresponds to a mental state of track position, task belief, and trial number (time). This structure is consistent with prior work showing that hippocampal population activity can exhibit disentangled codes with approximately orthogonal components across task variables and that context-related structure can remain preserved along dimensions largely orthogonal to ongoing drift [46, 123]. Importantly, by preserving local distances, angles, and directions, our latent space captures and reveals the quantitative relationships between different coded variables on the manifold in the learned representational space.

While our model of the cognitive map consists of three local directions, the vast majority of the mouse’s experienced mental state exists on two distinct task conditions (near, as in Figs. 5a,g, and far, as in Figs. 5b,h, rather than in between, as in portions of Fig. 5c), each of which predominantly occupies a 2-dimensional surface parameterized by position and trial (time). As such, for ease of analysis, we define the model of the mouse’s *experience* on the cognitive map as the two surfaces formed by averaging the coordinates of the data points embedded in the latent space according to discrete bins of position (3cm) and time (20 trials) (Fig. 5i, see Methods). This representation allows us to quantitatively project the maze coordinates through time onto these surfaces (grid lines in Fig. 5i).

### 8.2 Geometric decomposition and consolidation of single neuron place fields over trials

To complement our population-level decomposition of collective backshift along rotation and drift, we seek a similar decomposition for the entire place fields of individual neurons. Traditionally, single neuron backshift has been quantified using scalar measures such as the neuron center of mass [40, 41]. However, neuron tunings are often geometrically much more complex, exhibiting properties such as skew [27] and multiplex tuning [124], and drift [22, 23, 46], often violating the assumption that place fields can be thought of as uniquely coding for a specific position. Here we relax this assumption *via* a functional approach by treating each neuron’s activity across an entire trial as a continuous function, and decomposing changes in that function over time along a component of pure backshift/rotation and pure drift. To build intuition, if the place fields of all neurons simultaneously backshift by the same amount, then there is no drift, and the evolution of the representation is constrained to a phase along a 1-dimensional line (Fig. 6a). If instead the place fields backshift at different rates or change shape, there emerges an additional component of drift (Fig. 6b). Hence, while not fully indicative of the population-level representation, here we extract the single-neuron component of backshift.

To begin decomposing backshift and drift at the level of single neurons, we show the decoded activity of a single neuron across 50cm and ≈ 100 trials from the model manifold, and the corresponding raw data (Fig. 6c, top). At a specific trial *t*, we plot that neuron’s activity as a function of position, along with its activity at a later trial (Fig. 6c, middle blue), and its activity shifted in position indicating a pure rotation/backshift (Fig. 6, middle orange). We then decompose the place field change over trial (Fig. 6, bottom left) as a weighted sum of the pure rotational component (Fig. 6, bottom right, orange) and the residual which is the pure drift component (Fig. 6, bottom right, yellow). We further plot several examples of the model prediction and data for single neurons that largely contain a rotational component but backshift at different rates (Fig. 6d), thereby still resulting in a population-level drift. Additionally, while rotation/backshift is straightforward to characterize using statistics such as the center of activity for single place fields [41], our decomposition generalizes to arbitrarily complex fields. We plot additional examples of neurons that contain significant drift at the single-neuron level whether by changing field width, location, or relative spacing (Fig. 6e).

To study the statistics and evolution of single-neuron rotation, we compute the average rotational component for each neuron in animal A7 for the first to fourth quarter segments of all trials, and plot the distribution of that average rotation for all neurons (Fig. 6f). We observe a statistically significantly positive rotation (backshift) across all trial segments, and a narrowing of the variance of rotation for later trials. We report a statistically significantly positive average single-neuron rotation for all animals (Fig. 6g). Finally, to broadly characterize the phase-space of single neuron behavior, we plot the average rotational and drift component across all trials for each neuron and each animal with two example place fields (Fig. 6h), showing an almost complete lack of neurons that purely rotate or have negative rotation on average. In sum, we demonstrate a per-neuron decomposition of representational drift into components of pure rotation and pure drift, report significant per-neuron rotation across all animals, and define a phase-space of single neuron behavior.

